# Astrocytic-supplied cholesterol drives synaptic gene expression programs in developing neurons and downstream astrocytic transcriptional programs

**DOI:** 10.1101/2025.01.28.635252

**Authors:** Emilia Vartiainen, Dhara Liyanage, Illinca Mazureac, Rachel A. Battaglia, Matthew Tegtmeyer, He Jax Xu, Noora Räsänen, Julia Sealock, Steven McCarroll, Ralda Nehme, Olli Pietiläinen

## Abstract

Astrocytes participate in neuronal synaptic programs that are enriched for genetic associations in schizophrenia and autism spectrum disorders (ASD). To better understand how these co-regulated cellular programs are induced during early neuronal development, we studied astrocytes and iPSC-derived neurons in co-cultures and mono-cultures at 16 time points spanning 0.5 hours to 8 days. We found that upregulation in astrocytes of genes involved in cholesterol biosynthesis preceded the activation of synaptic gene programs in neurons and upregulation of the astrocytic *Nrxn1*. Neuronal knockdown of key cholesterol receptors led to downregulation of neuronal synaptic genes and induced a robust transcriptional response in the astrocytes, including further upregulation of *Nrxn1*. This suggests that astrocyte-supplied cholesterol drives these neuronal changes and that bi-directional signalling is occuring. The genes upregulated in neurons were enriched for deleterious variants in schizophrenia and neurodevelopmental disorders, suggesting that their pathogenic effect may be, in part, mediated by reduced buffering capacity for changes in the astrocyte cholesterol supply to neurons. These findings highlight the critical role of astrocyte-neuron interactions in psychiatric and neurodevelopmental disorders, particularly in relation to lipid metabolism and synaptic plasticity.

## Introduction

Genetic studies of rare and common variants in schizophrenia and autism spectrum disorders (ASD) have highlighted synaptic genes highly expressed by excitatory neurons^1–4^. These include a number of genes encoding glutamate signaling pathway members where protein truncating variants are associated with increased risk for these conditions^2,3^. New evidence emerging from single cell RNA sequencing (scRNA-seq) studies in primary brain tissue have found that in addition to neurons, glial cells, including microglia and astrocytes, also show cell type-specific changes in gene expression relevant to human disease^5,6^.

Many neuronal synaptic functions, from synapse development to maturation and maintenance, are regulated by neuron-surrounding astrocytes. Astrocytes control neuronal synaptic development and maturation by an array of secretion-based and contact dependent mechanisms^7–9^. It is well known that astrocytes in co-culture with neurons promote neuronal maturation *in vitro*. Previously, we found that astrocytes enhanced synaptic gene expression programs in human induced pluripotent stem cell (hiPSC)-derived mature neurons via contact-dependent mechanisms^10^. In turn, co-culture promoted increased expression of cell adhesion and cholesterol biosynthesis genes in astrocytes. More recently, similar and equally coordinated astrocyte-neuron gene expression programs have also been identified in human primary brain tissue ^6^. Concerted downregulation of neuronal synaptic gene expression coupled with decreased expression of synaptic adhesion and cholesterol biosynthesis genes in astrocytes have now been reported in both schizophrenia and ASD^5,6^. However, little is known about the nature of the underlying astrocyte-neuron co-regulation, and about how, and when, these pathways are triggered in astrocytes and neurons or the signals that promote them.

To bridge this knowledge gap, and understand how human genetic variation linked to psychiatric and neurodevelopmental disorders may alter neuron-astrocyte interactions, it is important to first understand the interplay of these cellular programs during development. In our earlier work, we focused on differentiated, mature neurons. Similarly, the human post mortem tissue-based studies captured mature cell states^5^. However, dissecting the timing and sequence of molecular changes during early development would not only afford a better understanding of disease trajectories, but may also illuminate more effective windows of therapeutic intervention to correct these altered trajectories.

HiPSC-based systems provide a uniquely powerful model to dissect and manipulate such early developmental events. Here, we studied gene expression programs activated by contact between hiPSC-derived early post-mitotic neurons and astrocytes, starting as early as 30 min and continuing through 8 days after co-culture. We discovered a transcriptional turning point at 24 hours post co-culture that overlapped with the synaptic neuron astrocyte program (SNAP) that was reported in the human brain^6^. This included upregulation of genes in cholesterol biosynthesis and synaptic adhesion molecules, such as *Nrxn1*, in astrocytes followed by increased synaptic gene expression in neurons with roles in psychiatric disorders. Knockdown of cholesterol uptake receptors, *LRP1* (with suggestive evidence of association to schizophrenia^49–51^) and *LDLR*, in neurons downregulated parts of the astrocyte-induced synaptic gene programs, and resulted in a highly correlated transcriptional response in astrocytes that involved upregulation of *Nrxn1* and monounsaturated fatty acid synthesizing *Scd2*. Blocking cholesterol uptake in neurons further resulted in increased expression of genes with deleterious coding variants in schizophrenia patients. Taken together, these findings provide novel insights into the mechanisms of co-regulation of early neuronal and astrocytic synaptic programs during maturation and highlight their role in neurodevelopmental disorders.

## Results

### Astrocytes undergo metabolic coupling with neurons within 24 hours of co-culture

To study genetic programs involved in astrocyte-mediated neuronal maturation, we designed a time-series experiment to measure gene expression in co-cultures across early development time (**Fig. 1a**). We differentiated hiPSCs towards an excitatory neuron fate using a deeply characterized Ngn2-based protocol in combination with forebrain patterning factors^11^. At 4 days post induction, the neural progenitor cells (NPCs) were co-cultured with mouse astrocytes derived from the cortex of neonatal mice^10^. We then used RNA-seq, resolved by species^10,12,13^, to study the transcriptomic profile of the co-cultured neurons and astrocytes at 16 time points, including several early time points (0.5, 1, 2, 3, 4, 5, and 6 hours) and daily intervals from 24 hours to 8 days, along with mono-cultures of both astrocytes and neurons harvested at the 48 hour time point (**Fig 1a**).

**Fig. 1:**
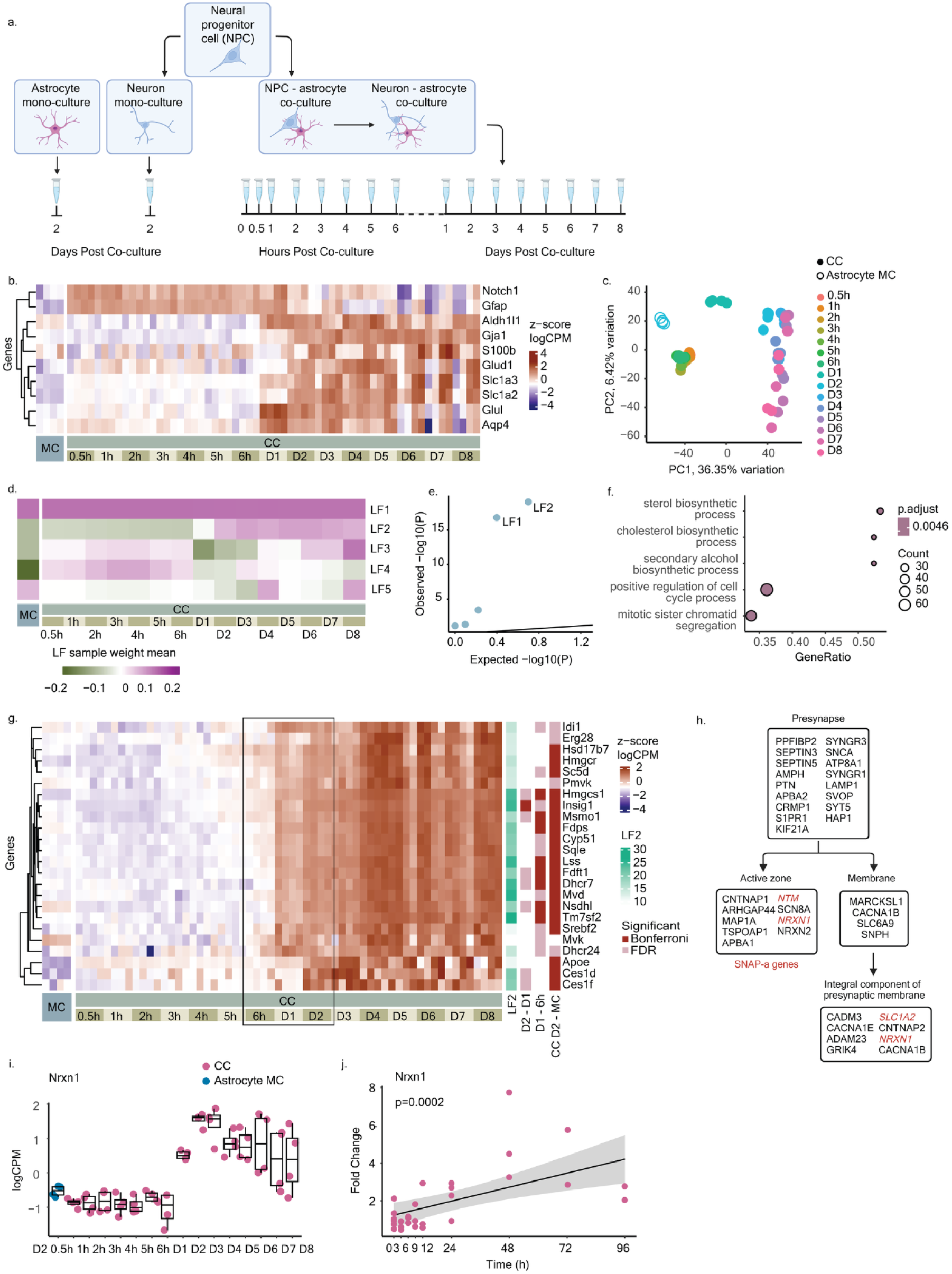
Astrocytes in contact with neuronal cells have increased expression of cholesterol biosynthesis genes. **a,** Outline of the experimental design. Human neural progenitor cells (NPCs) and mouse astrocytes were placed in mono-culture (left) or co-culture (right) (n = 4 biological replicates). Co-cultures were differentiated for up to 8 days. RNA-seq was conducted on mono- and co-cultures at the indicated time points post co-culture. Figure created with BioRender.com. **b,** A heatmap of standardized expression (z-score logCPM) of canonical astrocyte marker genes across all time points in the mono- and co-cultured astrocytes reveals changes in expression over time (n = 4 per time point). **c,** PCA of the astrocyte normalized read counts shows clustering of samples by time (colored, 0.5 h - day 8) and culture condition (fill, CC = co-culture, MC = mono-culture) across PC1 and PC2, explaining 36.35% and 6.42% of variation, respectively. **d,** A heatmap of mean factor values (sample weight mean) across time and culture condition from latent factor (LF) analysis of gene expression in astrocytes. **e,** A qq-plot of p-values from linear regression of LF values with time in co-culture reveals that LF2 has the strongest, significant association with time in co-culture. Expected p-value distribution shown by black line. **f,** Gene ratio of top enriched gene ontology (GO) terms from GSEA of the gene weights in LF2 show an enrichment for upregulated genes involved in cholesterol biosynthesis and lipid metabolism. **g,** Heatmap of standardized expression (z-score logCPM) of the sterol biosynthetic process genes (n = 24) show a distinct increase in expression between 6 hours and 2 days post co-culture. Colored annotations on the right of the heatmap show association of the gene with LF2 gene loading. Genes that are significantly differentially expressed between 2 days and 1 day in co-culture, 1 day and 6 hours in co-culture and between astrocytes in CC and MC at 2 days (p < 4.63908e-06 [Bonferroni cut-off] in red, FDR < 5% cut-off in pink). **h,** The presynaptic genes in LF2 include genes (n = 3) in the SNAP-a program in the human brain, which are highlighted in red. **i,** Boxplots of normalized expression (logCPM) for *Nrxn1* in astrocytes (n = 4 per time point). The boxplots’ limits show the upper- and lower quartile ranges, the center lines show the median values. The whiskers show the minimum and maximum values. **j,** Gene expression trends from qPCR experiment of *Nrxn1* at various time points after co-culture of human NPCs and mouse glia, along with the corresponding ANOVA test p-value.

After assigning the sequence reads to each species and confirming the cell identities of the astrocytes and the early postmitotic excitatory neurons (**Fig. 1b, Extended Data Fig. 1a, Extended Data Fig. 1b**), we set out to analyze gene expression patterns associated with the cocultures. We first, systematically explored the effect of co-culture on astrocyte gene expression using principal component analysis (PCA). Clustering of samples along the principal components PC1 and PC2 suggested notable transitions that took place immediately in response to co-culture (0.5 – 6 hours), at 6 – 24 hours, and 24 – 48 hours along PC1, implying distinct shifts in the astrocyte states in co-culture. These were followed by continuous, progressive changes in the transcriptome at later time points (day 2 – day 8) along PC2 (**Fig. 1c**), indicating that the astrocytes continuously adapted to the maturing neuronal cell states. The transcriptional transition at 24 hours of neuronal contact involved upregulated expression of characteristic astrocyte genes involved in glutamate recycling (*Glul, Glud1, Slc1a2, Slc1a3*), *Gaj1* and *Aqp4* (**Fig. 1b**), indicating metabolic coupling with neurons^14–18^.

### Contact with neuronal cells promptly induces expression of cholesterol biosynthesis genes in astrocytes

To interrogate gene expression changes in the astrocytes, we used a Bayesian approach for deducing hidden latent factors based on the co-variation in expression of many genes (Probabilistic Estimation of Expression Residuals, PEER)^6,19^. This allowed us to identify groups of similarly expressed genes, defined by gene loadings for each factor, and to study their expression levels simultaneously (the factor value) over the full course of the experiment. PCA indicated that the major sources of variance in the data were confined to only a few dimensions. To avoid overfitting of the data, we settled on 5 latent factors (LFs) in the analysis (**Fig. 1d**). Out of these factors, LF2 appeared to have a distinct set of underlying genes (loadings) that showed significant association with time in co-culture (**Fig. 1d,e, Extended Data Fig. 1c,e**).

Next, to identify biological functions of the astrocyte genes in LF2 that were associated with neuronal contact, we applied gene set enrichment analysis (GSEA)^20^ to the gene loadings of LF2. The analysis revealed several gene ontology (GO) terms related to cell proliferation that were enriched for downregulated genes in the astrocytes at 24 hours of co-culture, suggesting transition to a more mature astrocyte state (e.g., mitotic sister chromatid segregation (GO:0000070, adj. p-value = 0.00465, NES = -2.531). We observed a striking enrichment of upregulated genes in co-culture involved in cholesterol biosynthesis and lipid metabolism (GO:0006695, adj. p-value = 0.00465, NES = 2.614 and GO:0006629, adj. p-value = 0.00465, NES = 2.549) (**Fig. 1f, Extended Data Fig. 1g, 2a**) (**Supplemental Table S1**). These genes were first upregulated at 24 hours and their expression continued increasing with time plateauing at 48 hours. Out of the top 500 genes in LF2, 77 genes had functions in lipid metabolism, including members in the major enzymatic steps in production of cholesterol, such as the rate limiting HMG-CoA reductase (*Hmgcr*)^21^. The expression of cholesterol transporter *Apoe*^22^ and carboxylesterases *Ces1d* and *Ces1f* involved in the hydrolysis of cholesteryl esters to cholesterol ^23^ were upregulated at 48 hours following the upregulation of the biosynthesis pathway (at 24 hours). These gene expression changes are in line with astrocytes’ role as a major hub for cholesterol production in the brain^24^. We confirmed the change in expression levels of key genes through qPCR at several time points between 0 and 4 days of co-culture in both cell types. In astrocytes, we detected significant upregulation in genes involved in cholesterol biosynthesis, including *Srebf2* (*p* = 0.0205) (**Extended Data Fig. 4**). Collectively, these results indicated that astrocytic cholesterol metabolism was a key, early pathway almost immediately activated by interaction with neuronal cells and that it was associated with astrocyte maturation induced by neuronal presence.

### Induction of synaptic adhesion genes such as *Nrxn1* coincides with an increase in astrocytic cholesterol biosynthesis gene expression

Previously, we showed that astrocytic expression of synaptic adhesion genes was correlated with expression of synaptic programs in mature neurons^10^. We thus wondered whether LF2 involved increased expression of genes related to the role of astrocytes in synaptic functions. We performed an enrichment analysis of synaptic Gene Ontology (GO) terms in the SynGO database^25^ for the top 500 genes with the highest loading for LF2 and increased expression in co-culture. The analysis revealed a significant enrichment for astrocyte genes with annotated functions in neuronal presynapse, including the presynaptic active zone and membrane components (q-value = 8.19e-3, q-value = 2.75e-3, respectively) (**Fig 1h, Extended Data Fig. 1h**) (**Supplemental Table S2**). Among these was schizophrenia-associated *Nrxn1*^26–28^, whose expression closely followed the expression pattern of the cholesterol biosynthesis genes, suggesting they were co-regulated (**Fig. 1i**). The upregulation of synaptic adhesion genes, such as *Nrxn1* (**Fig. 1j**) and *Nlgn1*, was confirmed through qPCR (p = 0.000183, p = 0.00285, respectively).

Expression of cholesterol biosynthesis genes and the synaptic adhesion molecule *NRXN1* in astrocytes was also found to correlate with neuronal synaptic programs in the human brain as part of SNAP^6^. We found significant overlap of 6 genes in the astrocyte component of SNAP (SNAP-a) among the upregulated genes in LF2 (Fisher’s Exact Test p = 1.185e-4, OR = 10.26, total SNAP-a genes = 19), suggesting that these genes were induced by neuronal contact in the astrocytes. Besides *Nrxn1*, these genes included a synaptic adhesion gene (*Ntm*)^29^, genes encoding synaptic glutamate transporters (*Slc1a2* and *Slc1a3)*^30^, as well *Tmtc1*, fine-mapped to a common variant associated with schizophrenia ^1^, and *Rorb*, located in a locus linked to schizophrenia through common variant associations^1^. Overall, our findings suggested that the cell type interactions identified in the human brain were also present in our *in vitro* cell culture model, and that these genetic programs were initiated during early neuronal development.

Next, to explore whether genes expressed in astrocytes in contact with developing neurons were enriched with neurodevelopmental disorders, we performed a GSEA analysis for a list of genes with rare coding variants associated with neurodevelopmental disorders, ASD and schizophrenia^2,3,31–35^ as a gene set, for all LF2 gene loadings. We observed a significant enrichment (adjusted p = 0.0033, NES = 1.3519) for the genes with positive LF2 values (**Extended Data Fig. 3c**). These findings suggested that genes upregulated in astrocytes co-cultured with neurons were associated with neurodevelopmental disorders.

### Astrocyte LF2 is associated with increased expression of neuronal synaptic programs

Next, to understand the corresponding transcriptomic changes in developing neurons, we proceeded to characterize neuronal gene expression across co-culture time. Similarly to astrocytes, a PCA of the neuronal transcriptome separated neurons at time points recorded immediately after plating with astrocytes (0.5 – 6 hours) from the rest (days 1 – 8) (**Fig. 2a**). Neurons grown for 48 hours in mono-culture clustered separately from neurons in co-culture at the same time point, implicating the modular effect of astrocytes on neuronal maturation, in line with previous reports^10^.

**Fig. 2:**
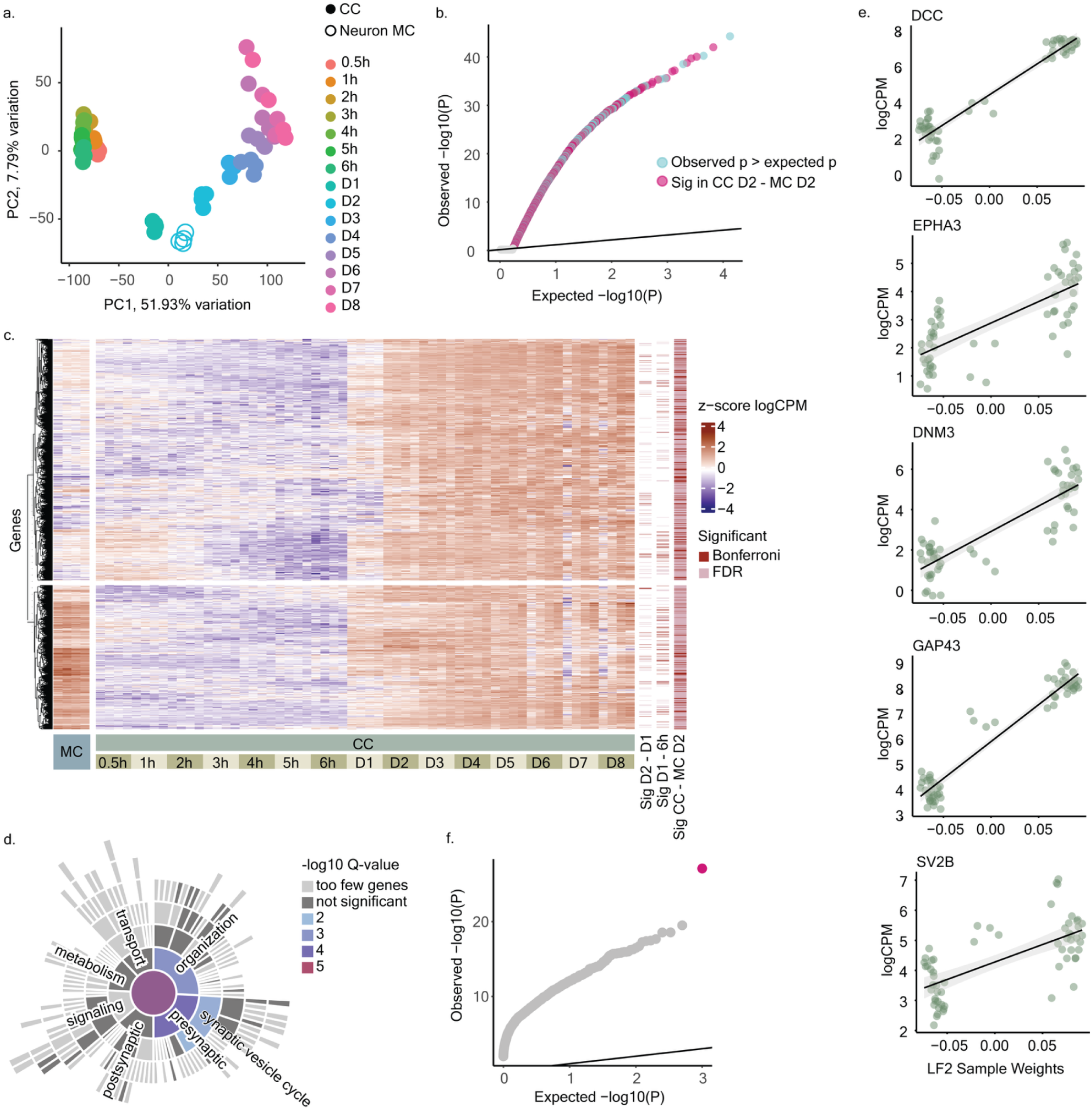
Astrocyte LF2 is associated with increased expression of neuronal synaptic programs. **a,** PCA of normalized read counts in neurons shows clustering of samples by time (colored, 0.5 h - day 8) and culture condition (fill, CC = co-culture, MC = mono-culture) across PC1 and PC2, explaining 51.93% and 7.79% of variation, respectively. **b,** A qq-plot of p-values from linear regression of neuronal gene expression values with LF2 sample weights. Astrocytic LF2 is associated with upregulation of 1,435 neuronal genes that have significant, positive association of expression changes in co-culture with the astrocytic LF2 gene programs. **c,** Heatmap of standardized expression (z-score logCPM) of the 1,435 upregulated genes associating with LF2 in co-culture. The genes cluster into two groups by hierarchical clustering. Colored annotations on the right of the heatmap indicate genes that were differentially expressed genes (p < 2.8991e-06 [Bonferroni]: red, FDR < 5%: pink) between: 1) neurons at day 2 and day 1 in CC, 2)neurons at day 1 to 6 hours in CC, and 3) neurons in CC and MC at day 2. **d,** SynGO sunburst plot showing enrichment of synaptic GO terms in the 842 genes with increased expression only in co-culture (Fig. 2c**, heatmap cluster 1**). The 842 genes were enriched in synaptic gene programs in synaptic vesicle cycle and synapse organization. **e,** A qq-plot of p-values from permutation analysis (n = 1,000 permutations) of variant burden in the SCHEMA consortium ^2^ data for genes associating with LF2 (n = 698). The p-values were calculated using Fisher’s exact test. The pink data point highlights enrichment of the gene set in question, the grey points demonstrate the permutation results. **f,** The neuronal genes upregulated exclusively in co-culture (Fig. 2c**, heatmap cluster 1**) included 3 SNAP-n genes: *DCC*, *EPHA3* and *DNM3;* the genes upregulated also in mono-culture (Fig. 2c**, heatmap cluster 2**) included 2 SNAP-n genes: *GAP-43* and *SVB2*. Scatter plots show neuronal normalized expression (logCPM) (y-axis) in the function of LF2 sample weights (x-axis). The black line is a fitted linear regression line with a 5% confidence interval.

Next, we aimed to identify neuronal genes whose expression correlated with the upregulated astrocyte gene programs in LF2. Linear regression revealed associations between astrocyte LF2 and global gene expression changes in maturing neurons, identifying 3,868 significant genes (adj. p < 0.05, beta > 0) (**Fig. 2b**). Among these, 1,435 genes were also differentially expressed (DE) between co-cultured and mono-cultured neurons, indicating co-culture-specific changes. Hierarchical clustering of these genes revealed two distinct patterns: 842 genes with high expression in co-culture at 24 hours and low expression in mono-culture, and 593 genes with the highest expression in mono-culture and the lowest in early co-culture (**Fig. 2c**). We confirmed these results through qPCR (**Extended Data Fig. 4**).

We previously showed that astrocytes induce expression of synaptic programs in mature neurons^10^. We thus investigated whether the 842 neuronal genes with increased expression at 24 hours were involved in synaptic gene programs annotated in SynGO. Interestingly, out of the 842 genes, 124 were associated with synaptic functions, with enrichments noted for biological processes such as the synaptic vesicle cycle (GO:0099504, p = 3.83e-4, n = 26 genes) and synapse organization (GO:0050808, p = 6.35e-5, n = 43 genes) (**Fig. 2d**) (**Supplemental Table S3**). Of these, 5 genes in the neuronal component of SNAP (SNAP-n) in the human brain were associated with LF2 (p = 0.0578, OR = 2.727, 95% CI = [0.780, 7.808]), 3 of which were upregulated only in the co-culture (**Fig. 2e**)^6^. These included genes associated with synaptic vesicle cycle (*DNMB*), synaptic adhesion (*EPHA3*) and postsynaptic density (*DCC*). Our findings suggested that these programs are initiated in neurons in part in response to astrocytes and correlate with expression of the astrocyte LF2 and an increase in astrocyte cholesterol gene expression.

Neuronal synaptic genes have been associated with common variants for schizophrenia, with SNAP genes showing reduced expression in individuals with the condition^6,10,36^. We wondered whether the genes with increasing expression in developing neurons had enrichment for damaging rare variants for schizophrenia. We calculated variant burden using data from the SCHEMA consortium^2^. Our analysis revealed that neuronal genes associated with LF2 (n = 698 found in the SCHEMA database) exhibited a significant burden of damaging rare variants in individuals with schizophrenia (p = 7.520e-28, OR = 1.218) (**Fig. 2f**). Out of the 698 genes, 482 (69%) had a higher frequency of damaging variants in schizophrenia cases than controls, suggesting that the mutation burden was not driven by any single gene. These results suggested that the neuron-astrocyte interactions in early neuron development might contribute to genetic risk for schizophrenia.

### Temporal changes implicate astrocytic cholesterol in driving expression of neuronal synaptic genes

To gain mechanistic insight into whether the initial transition in neuronal synaptic gene expression programs around 24 hours was driven by astrocyte gene programs or vice versa, we compared the temporal pattern of astrocytic expression of cholesterol synthesis genes and the neuronal expression of synaptic gene programs. To this end, we calculated the mean z-score of the expression values (logCPM) for the 124 synaptic genes that correlated with LF2 in neurons, as well as for the 52 upregulated astrocyte cholesterol genes involved in LF2 and astrocyte *Nrxn1* (**Fig. 3, Extended Data Fig. 5a)**. A temporal comparison of the standardized expression suggested that an increase in astrocyte cholesterol pathway genes began at 4-hours of co-culture, and therefore preceded the increase in expression in neuronal synaptic genes and astrocytic *Nrxn1* (**Extended Data Fig. 5a**) at 6 hours of co-culture. This sequential activation suggested that the upregulation of neuronal synaptic genes and astrocyte *Nrxn1* expression were driven at least in part by astrocyte-supplied cholesterol.

**Fig. 3:**
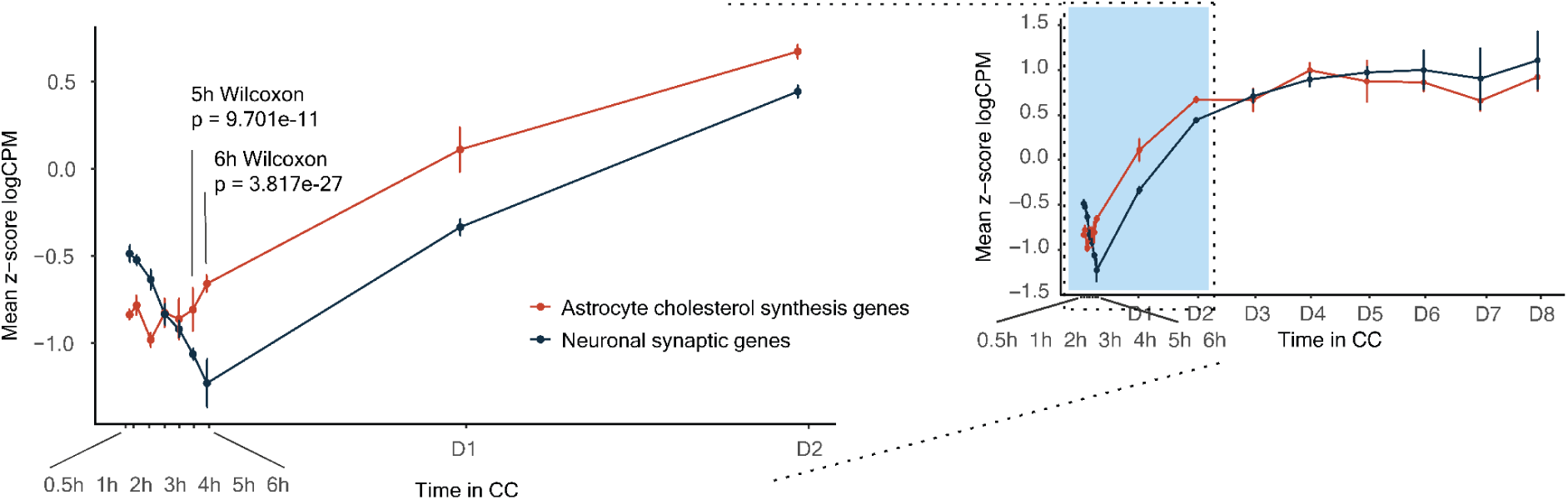
Temporal changes implicate astrocyte cholesterol in driving expression of neuronal synaptic genes. Comparison of the temporal pattern of expression of astrocyte cholesterol synthesis genes (n = 52) and the neuronal synaptic genes (n = 124). A plot of mean standardized expression (z-score logCPM) in the function of time up to day 8 (right) and an enlarged view of early time points (left). A two-sided Wilcoxon rank-sum test was used to calculate the statistical difference between the mean z-score logCPM values of astrocytic cholesterol genes and synaptic genes in neurons. The upregulation of cholesterol genes appeared to initiate before the upregulation of neuronal synaptic genes, suggesting that the neuronal synaptic gene expression was driven by astrocyte supplied cholesterol.

### Neuronal cholesterol receptors mediate pro-synaptic effects of astrocytes

The rapid increase of expression of cholesterol synthesis genes and *Apoe* in astrocytes led us to hypothesize that the subsequent induction of expression of synaptic programs in neurons was driven by astrocyte-supplied cholesterol. To test this hypothesis, we targeted two key cholesterol uptake receptors that bind *APOE* in neurons: *LDLR* and *LRP1*^37,38^. We employed CRISPR interference (CRISPRi)^39^ to knock down the gene expression of *LDLR* and *LRP1.* We designed three different guide RNAs (gRNAs) to knock down each gene, along with three non-targeting (NT) guides, which were introduced at the hiPSC stage through lentiviral transduction. The hiPSCs were then induced to generate NPCs, which were then co-cultured with astrocytes for 48 hours (**Fig. 3**), and profiled using RNA-seq to study the impact of the knockdown on neurons and astrocytes (**Fig. 4a**).

**Fig. 4:**
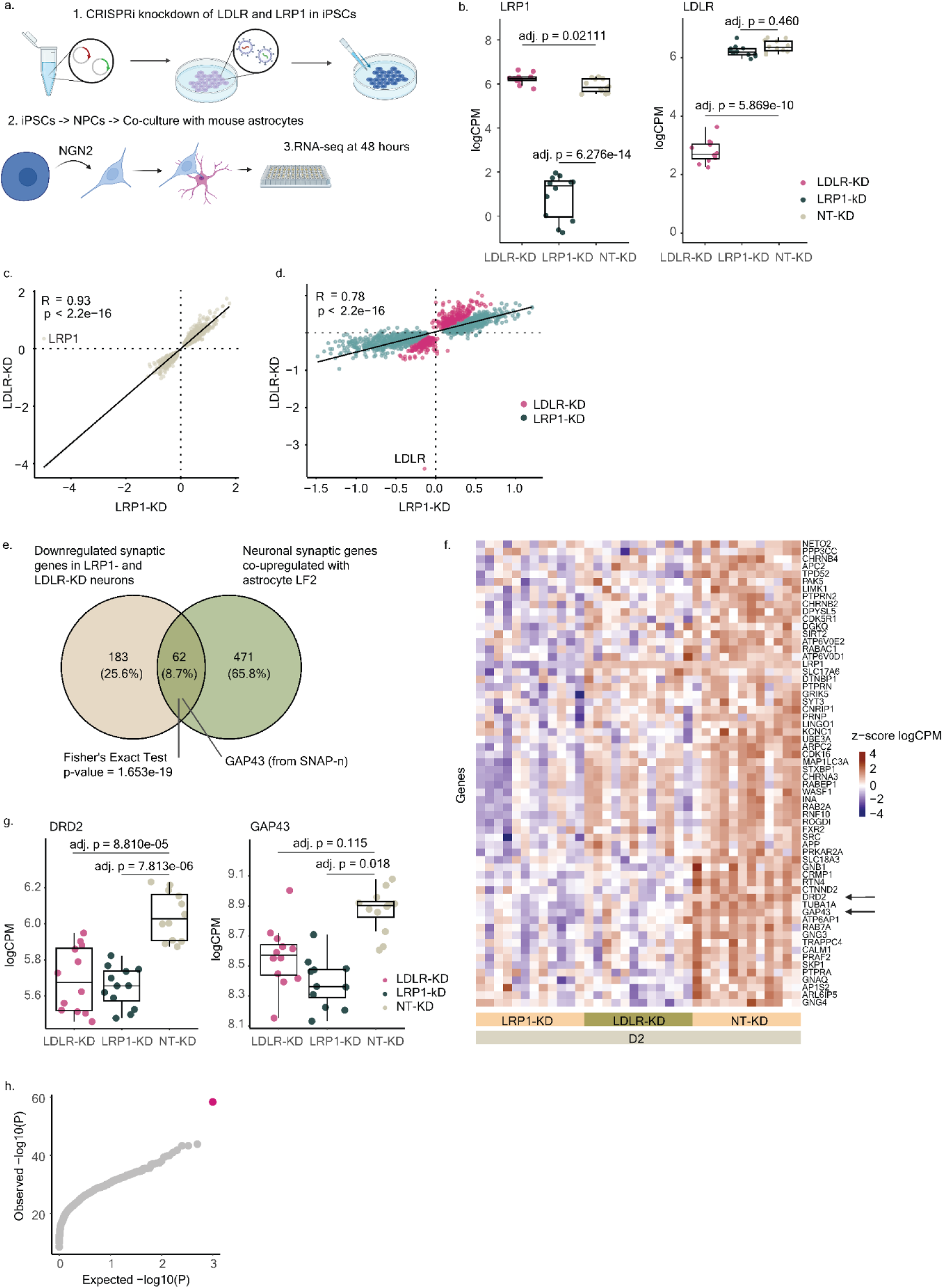
Neuronal cholesterol receptors mediate pro-synaptic effects of astrocytes. **a,** Experimental overview of the CRISPRi experiment. We used three different gRNAs to knock down two cholesterol receptor genes in neurons, *LRP1* and *LDLR*, along with three non-targeting (NT) guides to serve as control samples. The knockdown NPCs were plated with mouse astrocytes and cultured for 48 hours, after which they were harvested for RNA-seq. **b,** Boxplots of normalized expression (logCPM) of *LRP1* (left) and *LDLR* (right) show effective knockdown (KD) of the target genes by CRISPRi. The p-values were calculated using DREAM linear mixed models. **c-d,** Scatter plots of logCPM values from the differential expression (DE) analysis of the KD samples. The diagonal (y = x) is indicated with a black line. Correlation coefficients (R) and p-values were computed using Pearson correlation where LRP1 KD is teal and LDLR KD is pink. The DE analysis yielded similar results; the fold changes of the 1,774 shared significantly DE genes (**c**), as well as 1,858 unique significantly DE genes (**d**) were both highly correlated (R = 0.93 and p < 2.2e-16, R = 0.78 and p < 2.2e-16, respectively). **e,** A venn diagram of the neuronal synaptic genes downregulated in *LRP1*- and *LDLR*-KD neurons (n = 245, shown in beige) and neuronal synaptic genes upregulated with astrocyte LF2 (n = 533, shown in green) have an intersection of 62 genes (Fisher’s exact test OR = 4.637, p = 1.653e-19). **f,** Heatmap of standardized expression (z-score logCPM) of the 62 synaptic genes show a similar downregulation in both *LRP1*-KD and *LDLR*-KD against NT-KD samples. **g,** Boxplots of normalized expression (logCPM) of two SNAP-n genes, *DRD2* and *GAP43*, in *LRP1*-KD, *LDLR*-KD, and NT-KD neurons. The p-values were calculated using linear mixed models in the DREAM software package. The boxplots’ limits show the upper- and lower quartile ranges, the center lines show the median values. The whiskers show the minimum and maximum values. **h,** A qq-plot of p-values from permutation analysis (n = 1,000 permutations) of variant burden in the SCHEMA consortium^2^ data for upregulated genes in *LRP1*- and *LDLR*-KD neurons (n = 1,713). The p-values were calculated using Fisher’s exact test. The pink data point highlights enrichment of the gene set in question, the grey points demonstrate the permutation results.

We first examined the knockdown efficacy of each gRNA. All three gRNAs for *LDLR* and *LRP1* resulted in a significant, several-fold reduction of the target gene expression in *LDLR*-KD and *LRP1*-KD lines compared to the NT-KD lines (**Fig. 4b**). *LDLR*-KD resulted in slight upregulation of *LRP1* (adjusted p = 0.0211, logFC = 0.354), while *LRP1*-KD had no effect on the *LDLR* expression (adjusted p = 0.4602, logFC = -0.140), suggesting that the reduction in the level of one receptor was not substantially compensated by overexpression of the other receptor in the maturing neurons.

To identify the genes impacted by the knockdown of neuronal cholesterol receptors, we conducted differential expression (DE) analysis between *LDLR*-KD and *LRP1*-KD versus NT-KD samples (**Extended Data Fig. 6b)** (**Supplemental Table S6, S7**). The reduced expression of either of the cholesterol receptor genes appeared to markedly influence the cell state of the neurons, as demonstrated by thousands of DE genes (DEGs). Overall, the *LRP1*-KD was associated with more DEGs (FDR < 5%) than the *LDLR*-KD (3,165 vs. 2,231 DE genes, respectively), which were evenly divided between up- and downregulated genes (**Extended Data Fig. 6c**). These results confirmed a central role for cholesterol in the interaction. Given that all guides resulted in similar reduction of the target gene expression, we observed no dosage effect for the DEGs **(Extended Data Fig. 6d**).

Both LDLR and LRP1 bind Apoe containing lipoproteins, which are major transporters of astrocyte-supplied cholesterol in neurons^22^. To decipher whether the observed KD-associated differences in gene expression were related to changes in Apoe uptake in the neurons, we compared the fold changes of the DEGs in both conditions. Remarkably, 49% of DEGs (1,774 out of 3,632) in either KD experiment had significantly different expression in both KD-experiments with close to identical effect sizes (Pearson correlation R = 0.93) (**Fig. 4c**). Moreover, out of the rest of the 1,858 genes that were significantly DE in KD of one – but not the other – cholesterol receptor, all were changed to the same direction with highly similar effect sizes (Pearson correlation R = 0.78) (**Fig. 4d**). The striking similarity of the KDs suggested that the gene expression changes were likely due to reduced uptake of Apoe by the neurons caused by either of the receptors.

To identify biological functions affected by the KD of neuronal cholesterol receptors, we studied GO-term overrepresentation of genes changed in either of the conditions (**Supplemental Table S8, S9**). The downregulated genes were enriched for functions related to protein homeostasis (GO:0030162, adjusted p = 0.0344, NES = -1.2711) and energy metabolism (GO:0015980, adjusted p = 0.0074, NES = -1.367) (**Extended Data Fig. 7a**), while genes upregulated by the knockdown were enriched for several terms indicative of RNA localization (GO:0006403, adjusted p = 0.0034, NES = 1.510) and mRNA transport in cells (GO:0051028, adjusted p = 0.0078, NES = 1.505) (**Extended Data Fig. 7b**). Together the enrichments broadly suggested changes in cell metabolism and RNA homeostasis in the cells.

We then asked if the astrocyte-induced synaptic gene expression programs in neurons were regulated by astrocyte-supplied cholesterol. We reasoned that such cholesterol-induced genes would be downregulated in the KD-neurons. KD of the receptors reduced the expression of 245 genes with synaptic annotations in SynGO. Out of these, the expression of 62 genes were upregulated in astrocyte LF2 (Fisher’s exact test OR = 4.637, p = 1.653e-19) (**Fig. 4e**), suggesting that they were induced in neurons by the presence of astrocytes in a cholesterol-dependent manner. The 62 synaptic genes had the same direction of effect in *LDLR*-KD and *LRP1*-KD neurons; 20 were significantly changed in both, 37 genes were significant only in LRP1-KD, and 5 in LDLR-KD. We identified one SNAP-n gene, *Growth Associated Protein 43* (*GAP43*), that was downregulated in both of the KD neurons (**Fig. 4g**), suggesting that the expression of *GAP43* was regulated by cholesterol uptake of neurons. Another downregulated gene was *dopamine receptor D2* (*DRD2*) which is linked to schizophrenia through common variant associations^1^. *DRD2* is a leading target in antipsychotic drugs, treating neuropsychiatric disorders, bipolar disorder, schizophrenia and Parkinson’s disease^40^.

Earlier we showed that neuronal genes with increasing expression over time were enriched for damaging rare variants associated with schizophrenia. Next, we investigated whether genes altered in the KD-neurons carried a similar burden of these variants. As before, we utilized data from the SCHEMA consortium^2^ to extract the number of carriers of protein-truncating variants (PTVs) and missense variants. For the analysis, we grouped genes upregulated by *LDLR-KD* and *LRP1-KD* together and similarly grouped genes downregulated by *LDLR-KD* and *LRP1-KD*. Intriguingly, the upregulated genes (n = 1,712) demonstrated a highly significant enrichment (p = 4.688e-59, OR = 1.262) that remained robust through permutation analysis (**Fig. 4h, Extended Data Fig. 7c**). Out of the 1,712 genes, 926 had a higher damaging variant frequency in schizophrenia cases, suggesting that the mutation burden was not driven by a small group of genes. These findings suggested that the neuronal genes whose expression increased after blocking cholesterol uptake had a higher burden of deleterious variants in individuals with schizophrenia compared to controls.

### Co-cultured astrocytes respond to reduction in neuronal cholesterol uptake

To further explore the cholesterol-mediated bi-directional communication between astrocytes and neurons, we asked if neuronal KD of the cholesterol receptors influenced molecular programs in the accompanying astrocytes. PCA of the astrocytes co-cultured with *LRP1*-KD, *LDLR*-KD and NT-KD neurons separated the cholesterol receptor knockdowns from the NT-KD in PC1 and PC2 (**Fig. 5a**).

**Fig. 5:**
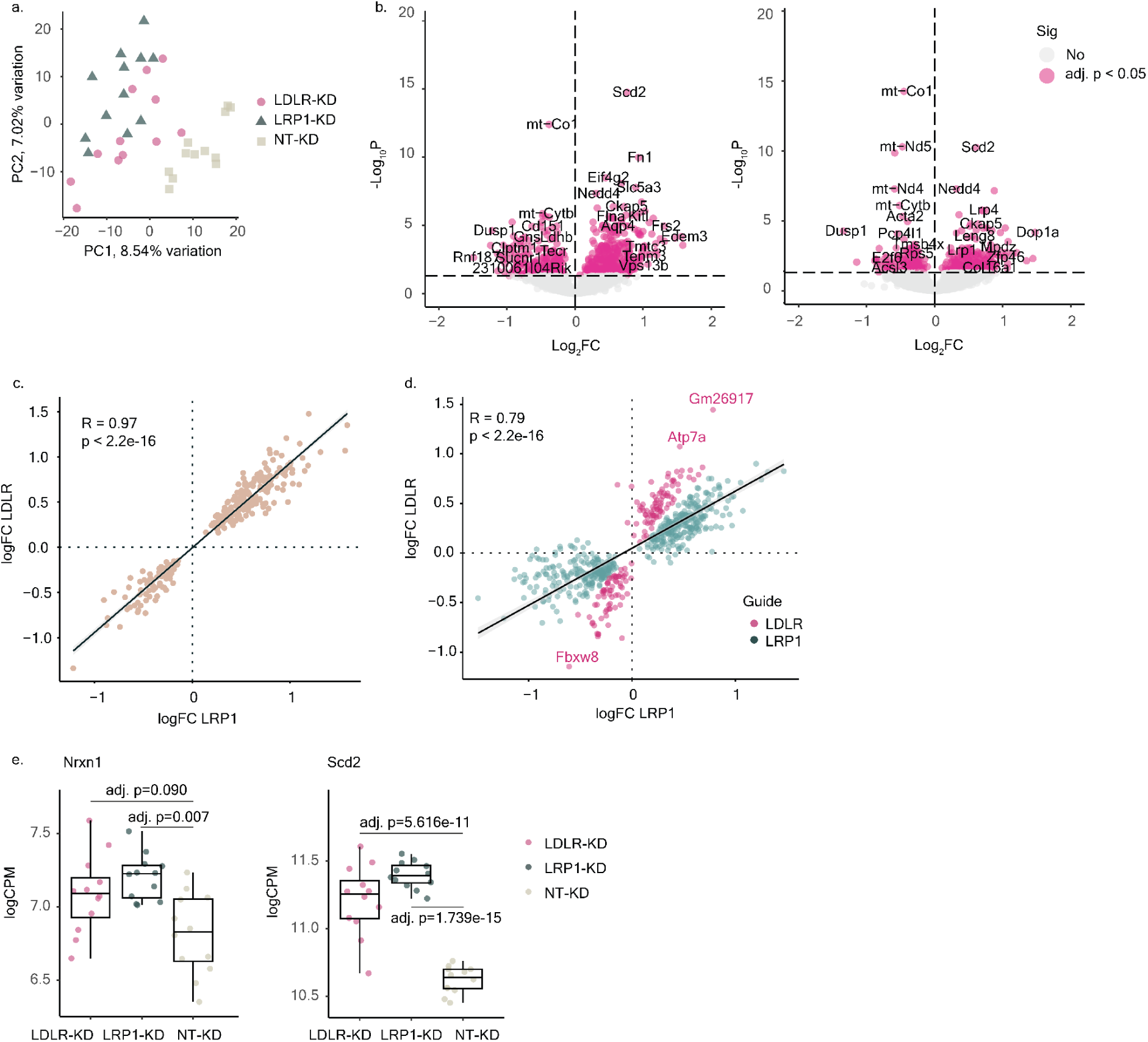
Co-cultured astrocytes respond to reduction in neuronal cholesterol receptor expression. **a,** PCA of the normalized read counts in astrocytes co-cultured with *LRP1*-KD, *LDLR*-KD and NT-KD neurons across PC1 and PC2 explaining 8.54% and 7.02% of variation, respectively. **b,** Volcano plots of differential expression analysis comparing astrocytes grown with *LRP1*-KD neurons or *LDLR*-KD neurons to the astrocytes co-cultured with NT-KD neurons. Pink fill highlights the significant (adjusted p-value < 0.05) genes. **c-d,** Scatter plots of logCPM values from the DE analysis of the astrocytes co-cultured with *LRP1*-KD, *LDLR*-KD and NT-KD neurons. Diagonal (y = x) is indicated by a black line. Correlation coefficients (R) and p-values were computed using Pearson correlation. **e,** Boxplots of normalized expression (logCPM) of *Nrxn1* and *Scd2* in astrocytes co-cultured with *LRP1*-KD, *LDLR*-KD and NT-KD neurons. The p-values are calculated using DREAM linear mixed models. The boxplots’ limits show the upper- and lower quartile ranges, the center lines show the median values. The whiskers show the minimum and maximum values.

A DE analysis further confirmed a highly identical response in the astrocyte transcriptome to the loss of either of the neuronal cholesterol receptors (**Fig. 5b**) (**Supplemental Table S10, S11**). Out of the 883 DEGs in astrocytes co-cultured with either *LDLR-*KD or *LRP1*-KD neurons, 97% had the same direction of effect in both treatments (R = 0.97, p < 2.2e-16), with 273 genes being significant (adj. p < 0.05) in both conditions (**Fig. 5c**). Of the 610 genes that were significantly DE in one but not the other condition, 591 had the same direction of effect and correlated fold-changes (R = 0.79, p < 2.2e-16) (**Fig. 5d**). As in neurons, *LRP1*-KD was associated with a slightly higher number of DE genes in astrocytes (n = 734) than *LDLR*-KD (n = 422). On average, astrocytes co-cultured with *LRP1-*KD neurons also displayed larger magnitudes of effects, with the mean of absolute logFC for shared DE genes being 0.5299 for *LRP1-*KD and 0.50980 for *LDLR*-KD. Taken together, these results showed that loss of either neuronal cholesterol receptor resulted in similar responses in the astrocytes, suggesting that this reaction was linked to a common change in the cellular environment caused by the loss of either receptor.

Notably, the neuronal KD of cholesterol receptors had no systematic, significant effect on the expression of cholesterol synthesis genes in the astrocytes (**Extended Data Fig. 8c**). Nevertheless, we noticed multiple genes for mitochondrial respiratory complexes (*mt-Nd1, mt-Nd2, mt-Nd4, mt-Nd5, mt-Nd6, mt-Co1, Mt-Cyb*) encoded in the mitochondrial genome were among the most significantly downregulated genes. This potentially indicated reduced energy demand in the astrocytes as a response to the neuronal loss of the key cholesterol receptors. To investigate if other biological pathways were implicated by astrocytic DEGs, we analyzed their enrichment for GO-terms (**Supplemental Table S12**). No specific biological functions were enriched for the downregulated genes beyond the large number of mitochondrial genes. For upregulated genes, on the other hand, the analysis highlighted several terms related to functions in neurite projection development (adj. p = 0.00756, NES = 2.429) and synapse organization regulation (adj. p = 0.03216, NES = 2.1572) (**Extended Data Fig. 9a**), potentially underlining an astrocyte reaction to impaired neuronal projection due to reduced cholesterol uptake.

*Nrxn1*, whose expression was correlated with the cholesterol biosynthesis genes in wildtype co-cultured astrocytes (**Fig. 1i**), was significantly upregulated in the astrocytes co-cultured with *LRP1*-KD neurons, suggesting it was regulated by cholesterol uptake in neurons (**Fig. 5e**). The single most significantly upregulated gene in the astrocytes in co-culture with *LDLR*-KD and *LRP1-*KD neurons was *Scd2* (p = 5.616e-11, logFC = 0.609 & p = 1.739e-15, logFC = 0.767, respectively) (**Fig. 5e**). Scd2 is the rate limiting enzyme in the synthesis of monounsaturated fatty acids (MUFAs), which are major constituents of the cellular membranes and bioactive lipids^41–43^. Taken together, these results underline a dynamic bi-directional interaction between astrocytes and neurons mediated by cholesterol to regulate neuronal maturation and function.

We next wondered whether any of these disease-associated genes enriched in LF2 had altered expression in astrocytes co-cultured with the KD-neurons. Indeed, *Nrxn1*^3,31,32^, *Kif1a*^33,34^*, Ank2*^3,31,35^ and *Map1a*^3^ had significantly (adjusted p < 0.05) upregulated expression in astrocytes co-cultured with KD-neurons (**Table 1**).

**Table 1:** Test statistics in astrocytes co-cultured with KD-neurons for disease-associated genes enriched in LF2. Values are from DE analysis comparing *LRP1-*KD and *LDLR*-KD neurons to NT-KD neurons. Statistically significant upregulation (adjusted p-value < 0.05) is indicated with an asterisk (*).

|  | *LRP1*-KD |  | *LDLR*-KD |  |
| --- | --- | --- | --- | --- |
| **Genes** | **adjusted p** | **logFC** | **adjusted p** | **logFC** |
| *Nrxn1* | 7.016e-03* | 0.384 | 0.090 | 0.292 |
| *Kif1a* | 1.843e-05* | 0.391 | 5.623e-03* | 0.283 |
| *Ank2* | 5.728e-06* | 0.683 | 1.682e-06* | 0.762 |
| *Map1a* | 0.177 | 0.446 | 3.841e-02* | 0.691 |

We next tested the up- and downregulated genes for enrichment of rare damaging variants for schizophrenia. Interestingly, as in the KD-neurons, the upregulated genes (n = 501) demonstrated a significant enrichment (p = 6.637e-19, OR = 1.202) (**Extended Data Fig. 9c**). The mutation burden was not driven by a single gene, as 281 genes had OR > 1. These findings further suggested that both neurons and astrocytes may have a capacity for compensating for changes in cholesterol metabolism across the two cell types, and this buffering ability appears to be diminished by the presence of pathogenic mutations.

## Discussion

To better understand pathways and genetic programs underlying astrocyte-neuron interactions during early neuronal development and maturation, and whether they might be vulnerable in neurodevelopmental and psychiatric disorders, we analyzed gene expression profiles evoked in astrocytes and early post mitotic neurons in co-culture. The analysis revealed a major transcriptional change in both cell types around 24 hours after co-culture. The changes appeared to be driven by astrocytic upregulation of cholesterol biosynthesis genes and synaptic adhesion molecules, such as *Nrxn1*, coupled with neuronal upregulation of synaptic genes. A knockdown of key cholesterol receptors *LRP1* and *LDLR* in neurons reduced neuronal synaptic gene expression and resulted in further upregulation of astrocyte *Nrxn1*. Nrxn1 is a synaptic adhesion molecule that is highly expressed by astrocytes, and it is likely involved in mediating physical interactions with neurons^44^. Loss of *NRXN1* is associated with schizophrenia and neurodevelopmental conditions^28,45^. Our findings suggest that *Nrxn1* and astrocyte-supplied cholesterol to neurons are co-regulated. Astrocytes supply cholesterol to neurons. This then triggers a molecular cascade in neurons that drives synapse maturation. Our data further suggests that synapse maturation then subsequently induces the expression of adhesion molecules, including *Nrxn1*. This in turn results in increased adhesion of astrocytes to neurons to support their ongoing maturation. Depletion of cholesterol uptake in neurons could, therefore, potentially disrupt adhesive interactions with astrocytes and cause compensatory upregulation of synaptic adhesion genes, such as *Nrxn1*, in astrocytes^46,47^.

Strikingly, the co-regulated gene expression programs in astrocytes and neurons overlapped with the synaptic neuron astrocyte program (SNAP) described in primary human brain tissue, which is reduced in schizophrenia and ASD^6,5^. Our findings suggest that the expression of several of the SNAP genes, including genes in the astrocyte cholesterol pathway, become coupled early in development and that the expression of SNAP in the adult human brain is causally linked to astrocyte-neuron interaction.

We further found that the upregulated genes in early post mitotic neurons had a higher burden of deleterious variants in individuals with schizophrenia compared to controls in the SCHEMA study^2^. This suggests that part of the genetic liability for schizophrenia could be mediated by astrocytes via neuron interactions in early development. In line with this, we also found an enrichment for genes involved in neurodevelopmental disorders that were specifically upregulated in the astrocytes in co-culture^2,3,31–35^. These mechanisms may involve cholesterol and impaired adhesive interactions between the cell types. Our findings provide insights into the developmental component of schizophrenia^48^ and further highlight astrocytes in the pathology of neurodevelopmental disorders.

Cholesterol is locally synthesized in the brain; and although both neurons and astrocytes synthesize cholesterol, the majority comes from astrocytes^24^ and is taken up by LDLR and LRP1 receptors in neurons. Suggestive evidence from smaller genetic and functional studies appears to point to *LRP1* variants in neuropsychiatric conditions^49–51^. For normal receptor activity, LRP1 is proteolytically processed by FURIN^52^. Notably, common variants affecting FURIN expression have been associated with schizophrenia ^51,53^ and with major depressive disorder (MDD)^54^, further highlighting the role of LRP1 in pathology of psychiatric disorders. Besides psychiatric disorders, LRP1 has been linked with the pathology of neurodegeneration in Alzheimer’s disease (AD)^55^, further underlining its key role for normal neuronal functioning and maintenance. We observed identical gene expression profiles with *LRP1*-KD and *LDLR*-KD, indicating that the observed changes may be mediated by the common cholesterol carrying Apoe ligand. The downregulated programs included numerous synaptic genes, in line with previous physiological observations^57,59^. *LRP1*-KD appeared to have a stronger effect on the transcriptome, indicated by the higher number of significant DEGs and overall larger effect sizes than detected for *LDLR*-KD. This could be due to higher endocytosis rates of *LRP1*^60^.

Interestingly, we observed an enrichment of deleterious schizophrenia-associated variants in the genes upregulated by *LRP1*-KD and *LDLR*-KD. Counterintuitively, this suggests that neurons may have a broad buffering capacity to compensate for changes in cholesterol uptake, which could then be diminished in the cases with rare mutations associated with schizophrenia. The effect of deleterious variants on disease typically follows a decrease in expression of the gene. Hence, it is possible that the resulting decrease in the expression range caused by the variants can impair the buffering capacity of the neuron for external variation in astrocyte supplied cholesterol to mediate their pathogenic effects. Similar mechanisms have been previously suggested for female protective effect in ASD^61^.

## Limitations of the study

In this study we show that cholesterol biosynthesis and lipid metabolism are induced in astrocytes co-cultured with early post mitotic neurons. By knocking down key cholesterol receptors in neurons we observe a downregulation of synaptic genes. While we performed a comprehensive analysis of gene expression in early astrocyte-neuron interactions, it will be important to look at protein and metabolite levels as well in the future. To examine astrocyte-neuron interactions, we used the widely accepted cross-species co-culture, which facilitated cell type-specific analysis of bulk RNA-seq data. However, human astrocytes differ from their rodent counterparts in many ways, including size and function^62^. Thus, a fully human co-culture system may better capture the cellular processes occurring in the human brain. Finally, although we highlight gene programs associated with schizophrenia and other neuropsychiatric disorders, we do not model them. Examining these programs in genetic backgrounds relevant to these conditions will be essential to reveal the specific ways these programs are altered in a pathological setting.

## Materials and Methods

### Differentiation of hiPSCs into excitatory neurons and bulk RNA-sequencing

Neuronal differentiation of the WTC-11 cell line was carried out by combining *NGN2* programming with SMAD and WNT inhibition as described previously^11^. NGN2 was integrated into the genome through electroporation, along with a doxycycline inducible promoter. The stem cells were fed with N2 supplemented media (Thermo Fisher, 17502048) containing doxycycline (2 μg/mL) and patterning factors SB431542 (Tocris, 1614, 10 μM), XAV939 (Stemgent, 04-00046, 2 μM) and LDN-193189 (Stemgent, 04-0074, 100 nM) for 3 days. At 4 days post induction, neuronal mono-cultures were switched to Neurobasal media (Gibco, 21103049) containing B27 (Gibco, 17504044, 50X), doxycycline (2 μg/mL), brain-derived neurotrophic factor (BDNF), ciliary neurotrophic factor (CTNF), and glial cell-derived neurotrophic factor (GDNF) (R&D Systems 248-BD/CF, 257-33 NT/CF, and 212-GD/CF at 10 ng/mL each). Neuronal co-cultures were grown in the same media formulation but were plated on top of murine glial cells at day 4 (mouse strain https://www.jax.org/strain/100012). The glial cells were derived from early postnatal mouse brains as described previously^63^. The neuronal mono-cultures and co-cultures underwent bulk RNA sequencing at the Broad Genomics Platform according to the Smartseq2 workflow^64^.

### Preprocessing of the human and mouse reads

To identify cell-type-specific gene expression profiles, the fastq files of the bulk RNA-seq data were aligned to a mixed human-mouse reference genome (GRCh37/hg19 and GRCm38/mm10, GSE63269) using STAR^65^ (v.2.5). The neuronal samples had an average library size of 1.8 million reads, with a standard deviation of 0.53 million reads. The astrocyte samples had slightly smaller libraries, averaging 1.2 million reads with a higher standard deviation of 1.3 million reads.

As quality control, Illumina adapters and low-quality base bares were trimmed using Trimmomatic^66^ (v.0.36) and short reads (< 36 base pairs) were removed. Next the reads were annotated with GENCODE GTF annotation version 19 and a custom GTF file for the mixed genome (GSE63269) for the mixed genome. A raw counts matrix was generated using featureCounts^67^ by Rsubread (v.1.32). Genes with a total read count less than 10 were filtered using filterByExpr function in edgeR^68^ (v. 4.2.0).

Read counts were then normalized for library size (method = TMM) and transformed to log2 counts-per-million (logCPM) estimates using the voom function by limma^69^ (v. 3.60.3) package. The normalized logCPM gene expression matrices were used as input for all the following analyses.

### Latent factor analysis

A latent factor analysis was performed on combined dataset of the astrocytes grown in mono-culture for 48 hours, as well as astrocytes grown with neurons in the time series experiment, which was RNA-sequenced at 16 time points, including several early time points (0.5, 1, 2, 3, 4, 5, and 6 hours) and daily intervals from day 1 to day 8. The analysis was performed using PEER (v.1.0)^19^ with the default parameters.

Given the experimental design, the data was presumed to have only few major sources of variance. Therefore, to avoid overfitting, the number of latent factors to be determined was set to 5. To verify the number of latent factors, the analysis was replicated with 8, 10 and 15 latent factors. To verify our choice of latent factors, the data was examined by identifying different numbers of latent factors in the data, all of which replicated the same key findings. With both 5 and 15 latent factors, the sample and gene weights of LF2 correlated at 99%, demonstrating that the results were robust regardless of the number of factors used. For clarity, the results of the latent factor analysis were multiplied by -1.

For analyzing association between genes in latent factors and time in co-culture, alinear regression analysis was conducted using the lm() function in R. The analysis tested the sample weights of each latent factor against two variables; time and cell culture.

### Gene set enrichment analysis

Gene set enrichment analysis was performed using gseGO from the clusterProfiler^70^ package (v.4.12.0) in R with org.Mm.eg.db (v.3.19.1) for mouse gene sets and org.Hs.eg.db (v.3.19.1) for human gene sets. 10,000 permutations were computed. A minimum of 10 and maximum of 500 genes per category were set as limits to the analysis. Benjamini & Hochberg method was used to calculate the adjusted p-values, with p-value < 0.01 and adjusted p-value < 0.05 cutoffs.

GO term overrepresentation analysis was performed using the enrichGO function from the clusterProfiler^70^ package (v.4.12.0) with the same parameters as listed above. In addition, a custom gene list was used as the universe in the background for the analysis, consisting of all genes found in the RNA-seq data.

For the enrichment of synaptic annotations, the online portal of SynGO^25^ was used. The analysis was performed using a custom gene universe of all the genes included in the RNA-seq data.

### Differential expression analysis

Differential gene expression analysis was performed using the DREAM software^71^ by variancePartition (v.1.34.0) package. The DREAM analysis pipeline was followed and the advanced hypothesis testing to compare relevant coefficients was performed using a contrast matrix. The astrocyte reads and neuron reads were analyzed separately, however a similar model was used for both analyses. Information of culture conditions (cells grown in co-culture of astrocytes and neurons or mono-culture) and the time point of sequencing were combined into one parameter (culture_time) so that the model used in the differential gene expression analysis was ∼ 0 + culture_time. For the CRISPR edited neurons the model included the target gene of the gRNA and the gRNA as a random effect, ∼ 0 + target_gene + (1 | gRNA). In order to increase power for the analysis of astrocytes grown in co-culture with the CRISPR edited neurons, only the target gene of the gRNA was used in the model, ∼ 0 + target_gene.

### Rare variant burden analysis

Enrichment of rare deleterious variants for schizophrenia was performed using data from the SCHEMA consortium^2^. The number of carriers of protein-truncating variants (PTVs) and missense variants was extracted for various gene sets (neuron genes associating with LF2, *LRP1*-KD and *LDLR-*KD neuron up- and downregulated genes, astrocytes co-cultured with *LRP1*-KD and *LDLR-*KD neurons up- and downregulated genes). Fisher’s exact test was used to calculate p-values and odds ratios (ORs) for the enrichment. To validate the observed variant burden, permutation testing (n = 1,000) was performed using randomly generated gene sets matched to the original sets by expression level (±20%) and size.

### qPCR

qPCR was performed using the previously described human neuron mouse glia co-culture system. Cell Lysates were prepared from six-well plates at the following time points: 0, 3, 6, 9, 12, 24, 48, 72, 96 (hours post co-culture). RNA was isolated using the RNeasy Kit (Qiagen, 74106) and converted to cDNA using the iScript cDNA Synthesis Kit (Bio-Rad, 1708890). The samples were prepared using the iQ SYBR Green Master Mix (Bio-Rad, 1708880) and qRT-PCR was performed using the C1000 Thermal Cycler (Bio-Rad, 32987). RPL10 was used as a housekeeping gene in both species. The primer sequences are as follows (5’-3’):

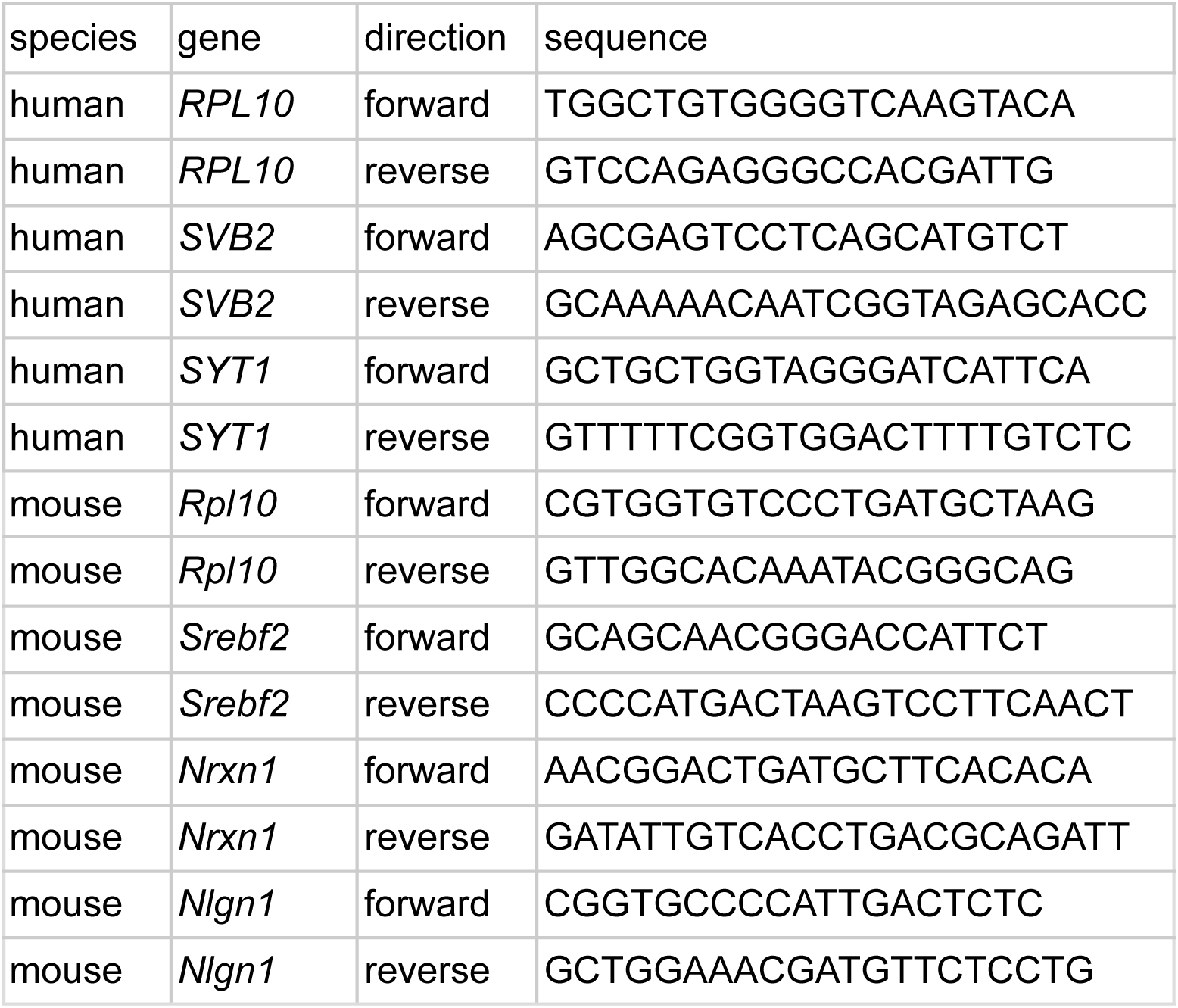

### CRISPRi gene editing

sgRNAs targeting the transcriptional start site of genes *LDLR* and *LRP1* were generated using the CRISpick technology^72,73^. Using the Golden Gate cloning protocol^74,75^, the sgRNAs were cloned into a CROPseq vector, packaged with *Trans*IT-293 reagent (Mirus Bio, MIR 2704) and packaging plasmids VSV-G and DVPR (Addgene, 12259, 12259). HEK239T cells (ATCC, CRL-3216) were transfected with the sgRNA plasmids. Lentivirus was concentrated from the cell media through the LENTI-X concentrator (Takara, 631231). The WTC11_TO-NGN2_dCas9-BFP-KRAB human iPSC line^76^ was plated onto Geltrex-coated plates with mTeSR1 media and incubated with the sgRNA lentivirus. After 48 hours, the iPSC culture media was fully switched to mTeSR1 supplemented with 1 ug/ml puromycin (Sigma Aldrich, P8833) to select for iPSCs that had uptaken the sgRNA.

## Acknowledgments

We thank members of the Nehme and the Pietilainen labs as well as Kris Dickson for helpful comments and discussion. The authors wish to acknowledge CSC - IT center for Science, Finland, for computational resources.

## Contributions

O.P. and R.N. conceived the work. E.V., O.P. and R.N. wrote the manuscript with input from all authors. D.L., I.M., R.B., J.X., and M.T. performed the experiments. E.V. processed and analyzed the data. R.N., and O.P. supervised the work and analyses. All authors edited and approved the final manuscript.

## Extended Data Figures

**Extended Data Fig. 1:**
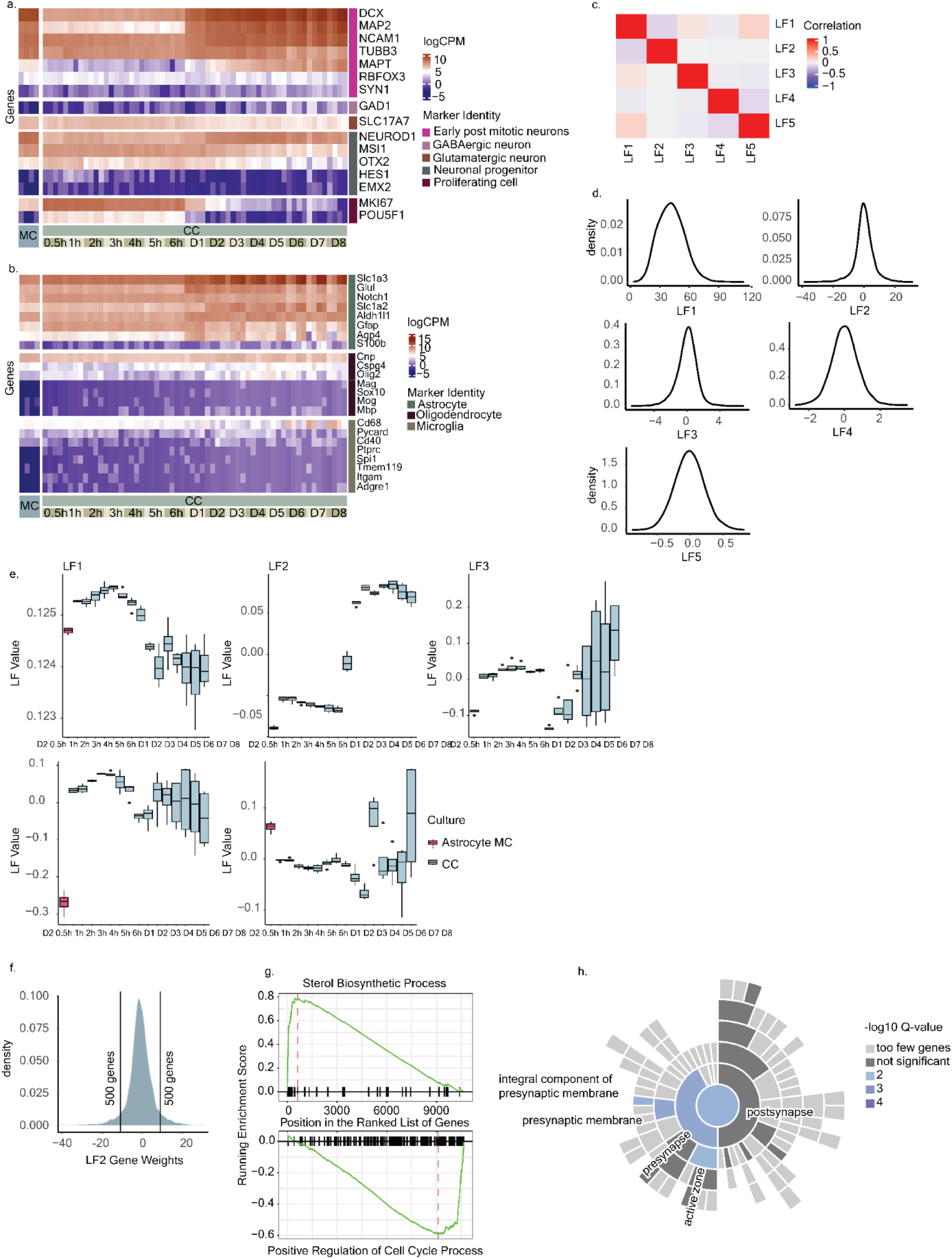
Characterization of neuronal and astrocyte samples, properties of the latent factors. **a,** Heatmap of normalized expression (logCPM) of canonical neuron marker genes in neuronal samples (n = 68). The genes are divided into clusters based on their marker identity, shown on the annotation on the right of the plot. Human cells in all cultures expressed genes characteristic for early neuronal identity. Neurons showed decreasing expression levels of neuronal progenitor marker genes and proliferative marker genes over time but had only low expression of mature neuron markers, in line with the expected early post-mitotic neuron identity. High levels of glutamatergic neuron marker *SLC17A7* was observed across all time points, while GABAergic neuron marker *GAD1* remained low, confirming that the neurons were predominantly excitatory. **b,** Heatmap of normalized expression (logCPM) of canonical astrocyte marker genes in astrocyte samples (n = 68). The genes are divided into clusters based on their marker identity, shown on the annotation on the right of the plot. Mouse cells in mono- and co-culture exhibited abundant expression of genes characteristic of astrocytes confirming the expected astrocyte identity of the cell population. **c,** Heatmap of Pearson correlation values of gene weights in each latent factor. **d,** Gene weight distributions for each latent factor. **e,** Boxplots of latent factor sample weights. The boxplots’ limits show the upper- and lower quartile ranges, the center lines show the median values. The whiskers show the minimum and maximum values. **f,** A density plot of the distribution of gene weights (n = 10848 genes) in LF2 were normally distributed with tails having the strongest positive (highest loadings) and negative (lowest loadings) association with the latent factor. **g,** Gene ratio of enrichment scores (ES) from GSEA of the gene weights in LF2 show an enrichment for upregulated (top) and downregulated (bottom) genes. The leading edge subset contributing most to the ES and top enrichment are marked with a dashed, red line. Genes in the respective GO terms are highlighted in the ranked order of LF2 gene weights by vertical lines on the x-axis. **h,** Sunburst plot displays enrichment of synaptic GO terms in the top 500 genes in LF2 from SynGO ^25^, showing that LF2 is enriched for presynaptic annotations in SynGO.

**Extended Data Fig. 2:**
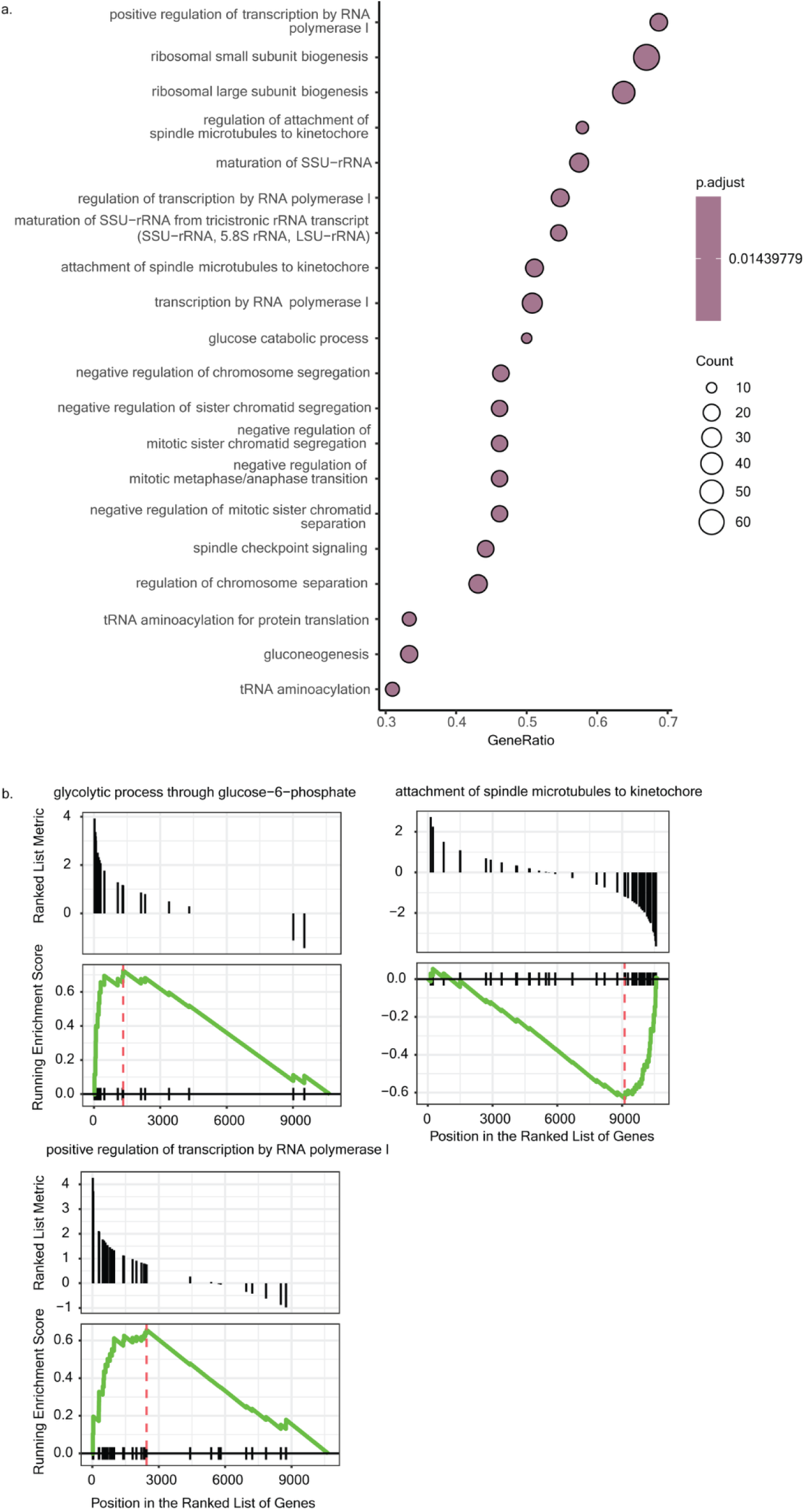
Properties of latent factor. **a,** Gene ratio of top enriched GO terms in LF3. **b,** Enrichment scores (ES) from GSEA of the gene weights in LF3. The leading edge subset contributing most to the ES and top enrichment are marked with a dashed, red line. Genes in the respective GO terms highlighted in the ranked order of LF2 gene weights indicated on the x-axis.

**Extended Data Fig. 3:**
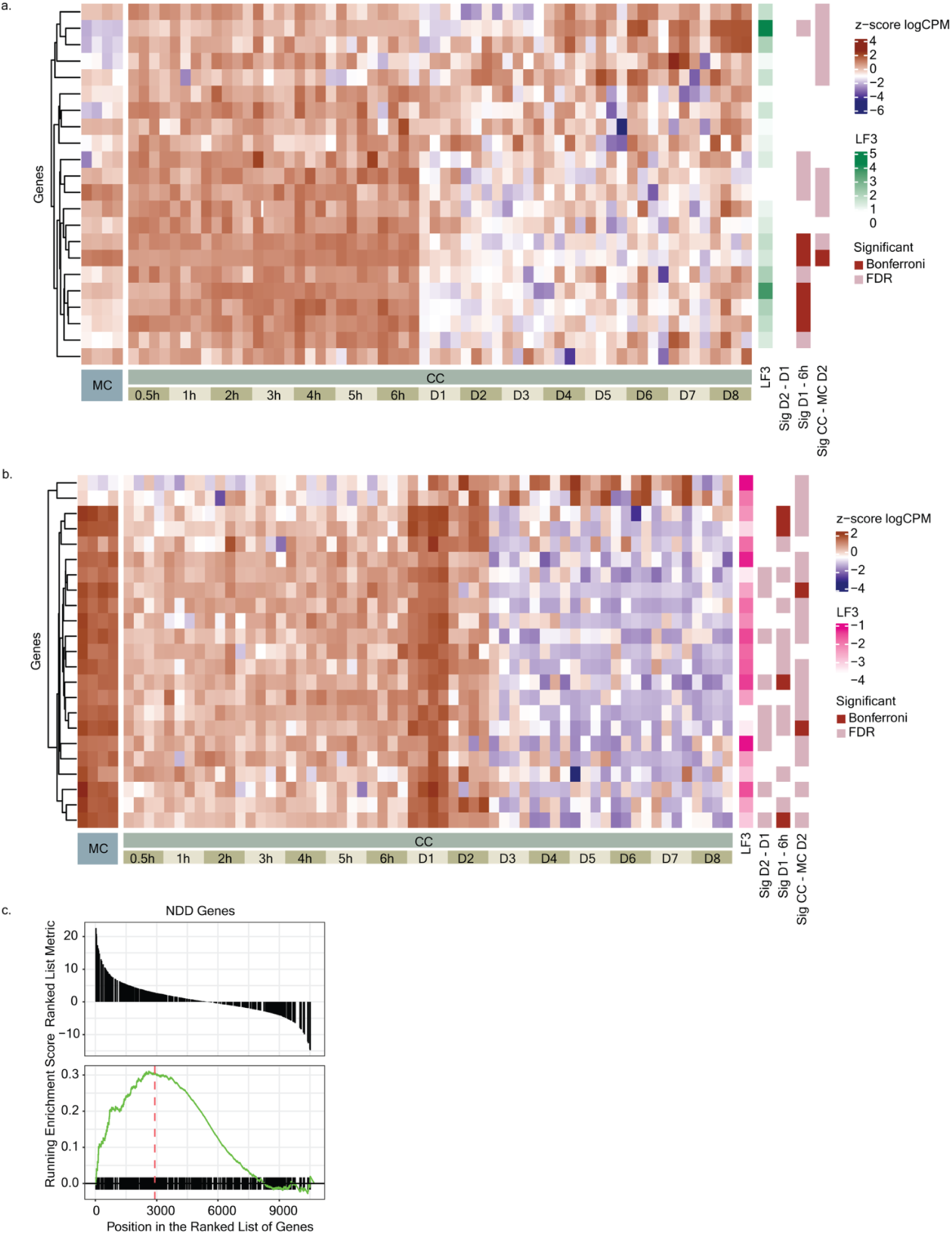
Properties of latent factor. **a,b,** Heatmap of standardized expression (z-score logCPM) of the positive regulation of transcription by RNA polymerase I genes (**c**) and attachment of spindle microtubules to kinetochore (**d**) in astrocytes (n = 68). Annotations on the right of the plot show association of the gene with LF2, significance from differential expression analysis for day 2 to day 1, day 1 to 6 hours and CC day 2 to MC. **c,** Enrichment scores (ES) from GSEA of neurodevelopmental disorder (NDD) genes as gene set, against LF2 gene weights. The leading edge subset contributing most to the ES and top enrichment are marked with a dashed, red line. NDD genes highlighted in the ranked order of LF2 gene weights indicated on the x-axis.

**Extended Data Fig. 4:**
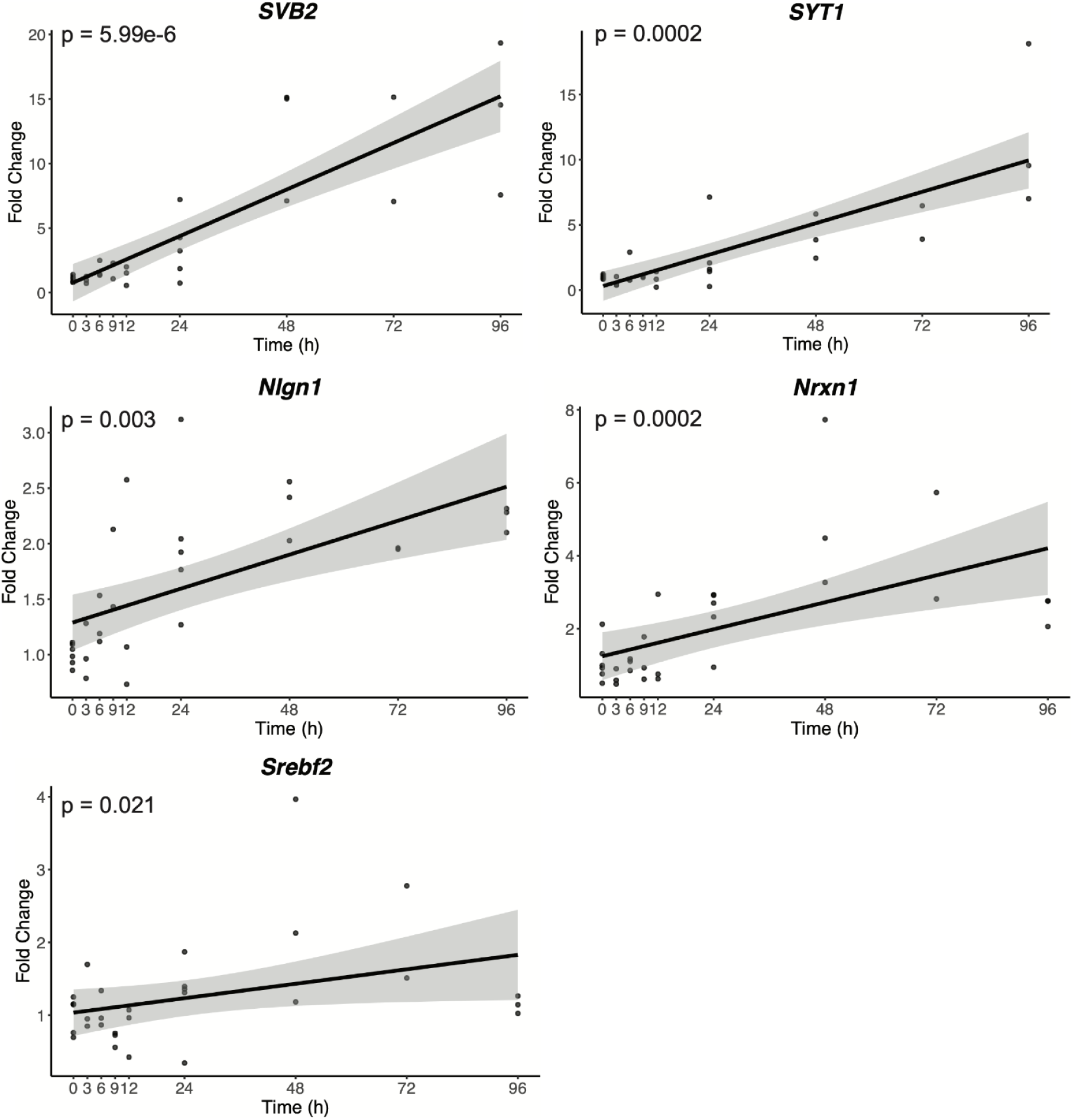
Validation of transcriptional changes in co-cultured neurons and astrocytes. Gene expression trends of selected genes at various time points after co-culture of human NPCs and mouse glia, along with the corresponding ANOVA test p-value. The human neurons showed significant upregulation of synaptic genes *SVB2* and *SYT1*. Thebon mouse glia showed significant upregulation of synaptic adhesion genes *Nrxn1* and *Nlgn1* as well as *Srebf2*, a gene implicated in cholesterol biosynthesis.

**Extended Data Fig. 5:**
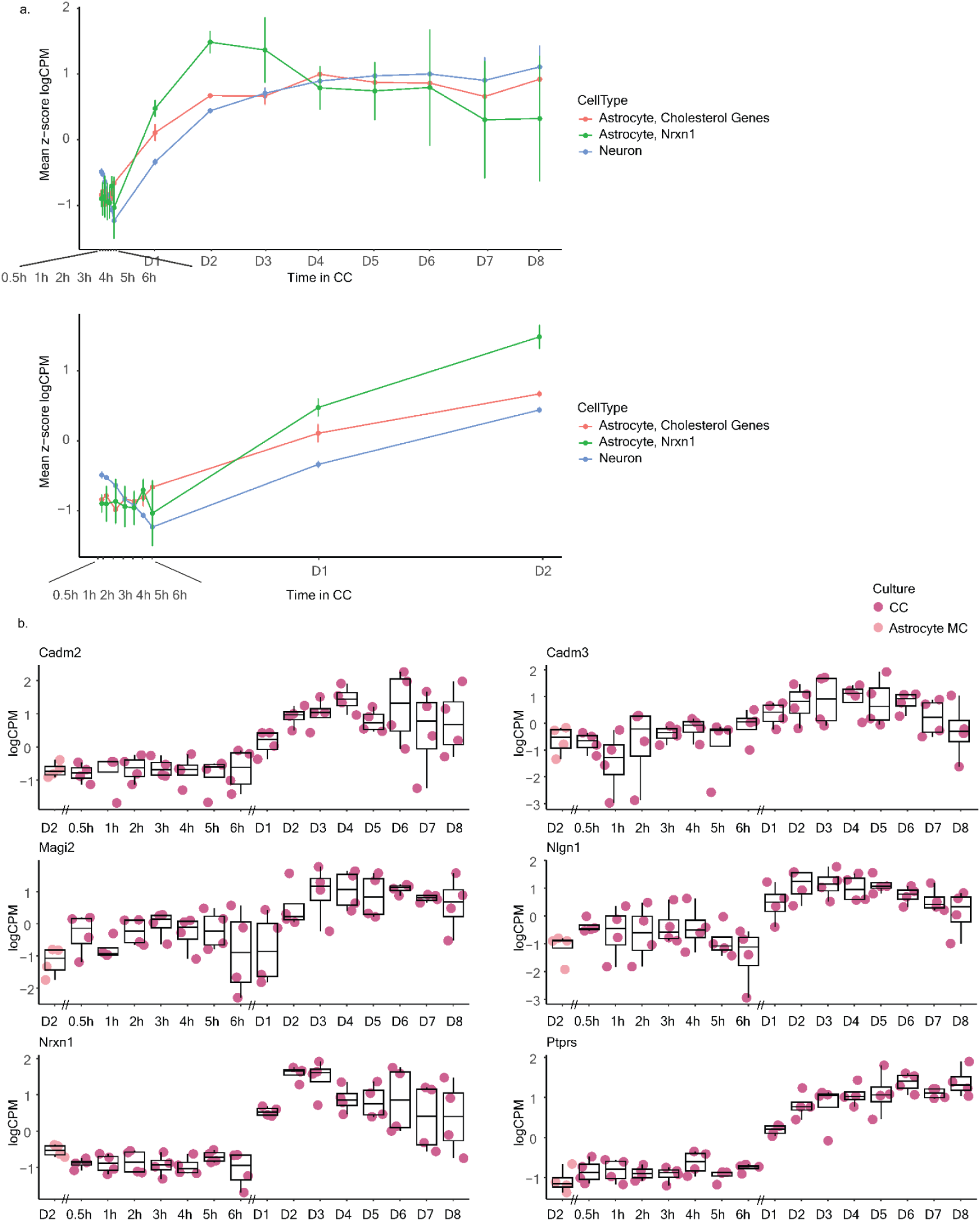
Timeline comparison of Nrxn1 expression with astrocyte cholesterol and neuronal synaptic genes. **a,** Comparison of temporal pattern of expression of *Nrxn1*, astrocyte cholesterol synthesis genes (n = 52) and the neuronal synaptic genes (n = 124). A plot of mean standardized expression (z-score logCPM) in the function of time up to day 8 (top) and an enlarged view of early time points (bottom). **b,** Boxplots of normalized expression (logCPM) for synaptic adhesion genes (n = 6) in astrocytes (n = 4 per time point). The boxplots’ limits show the upper- and lower quartile ranges, the center lines show the median values. The whiskers show the minimum and maximum values.

**Extended Data Fig. 6:**
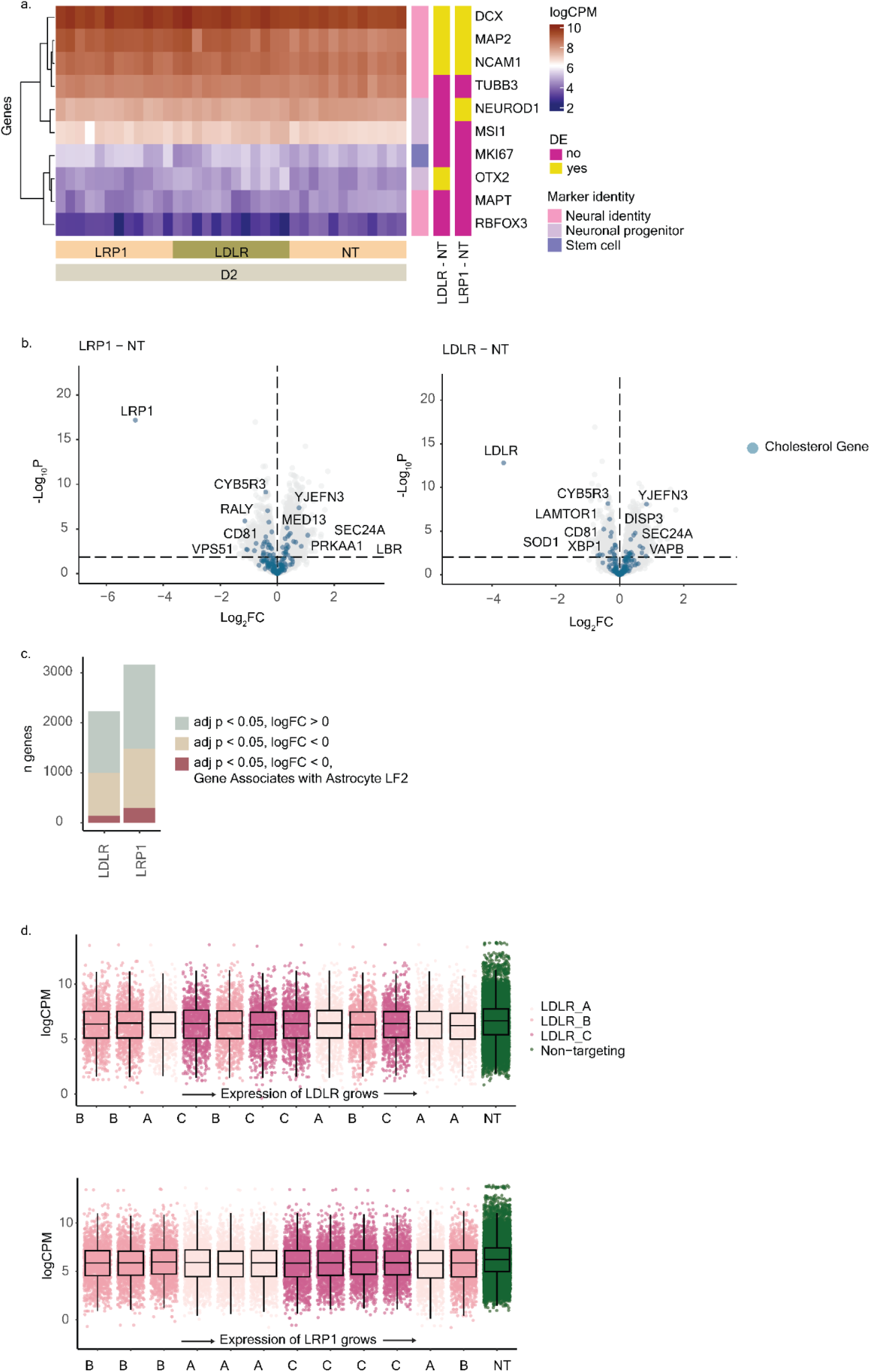
CRISPRi-KD neurons. **a,** Heatmap of normalized expression (logCPM) of canonical neuron marker genes in CRISPRi-edited neuronal samples (n = 36). Annotations on the right of the plot show marker gene identity and whether gene is significantly DE between in the knockout samples (adj. p < 0.05), obtained using DREAM linear mixed models. **b,** Volcano plots from differential expression analysis comparing *LRP1*-KD and *LDLR*-KD to NT-KD samples. Log2(FC) values on the x-axis, -log10(P) values on the y-axis. Horizontal dashed line shows a threshold for significance, adj. p = 0.05. Vertical dashed line shows log2(FC) = 0. Genes related to cholesterol metabolism are colored blue. **c,** Barplot showing the number of DEGs in the analysis on plot B. Turquoise color shows upregulated genes (adj. p < 0.05 & log2(FC) > 0), beige color shows downregulated genes (adj. p < 0.05 & log2(FC) < 0) and red color shows the number of downregulated genes that, in neurons, associate with astrocyte LF2 values. **d,** Box plots of normalized expression (logCPM) of downregulated genes (logFC < 0, adj.p < 0.05) in *LDLR* (upper plot) (n = 993 genes) and in *LRP1* (lower plot) (n = 1438 genes) all gRNA knockdown lines (A, B, C) and NT lines. x-axis is ordered by the target gene (*LDLR*, *LRP1*) expression, from lowest to highest.

**Extended Data Fig. 7:**
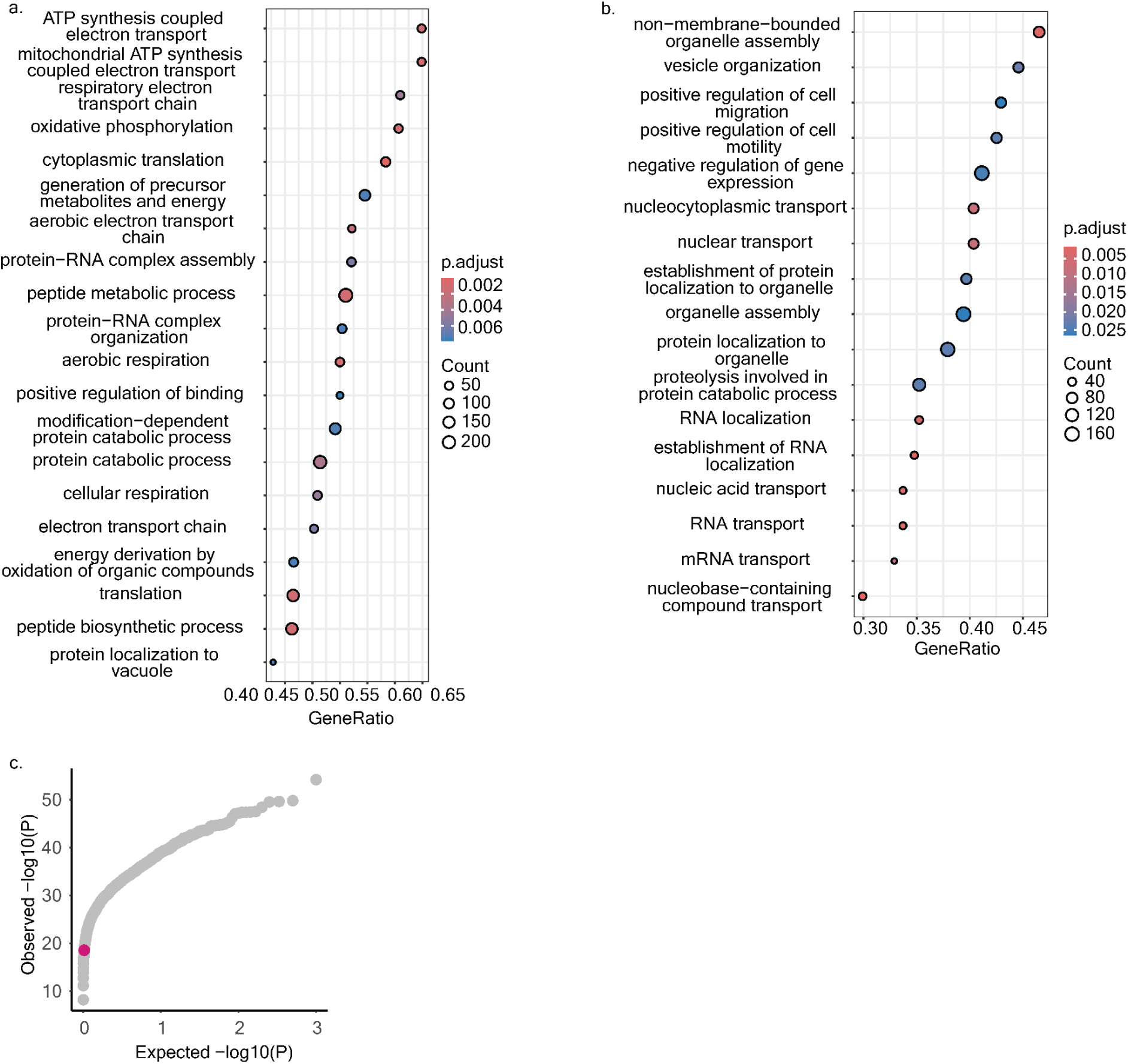
CRISPRi-KD neurons GO terms and SCHEMA variant burden analysis. **a,** Gene ratio of top enriched GO terms in downregulated genes in *LRP1*- and *LDLR*-KD neurons. **b,** Gene ratio of top enriched GO terms in upregulated genes in LRP1- and *LDLR*-KD neurons. **c,** A qq-plot of p-values from permutation analysis (n = 1,000 permutations) of variant burden in the SCHEMA consortium ^2^ data for downregulated genes in *LRP1*- and *LDLR*-KD neurons (n = 1,618 genes). The p-values were calculated using Fisher’s exact test. The pink data point highlights enrichment of the gene set in question, the grey points demonstrate the permutation results.

**Extended Data Fig. 8:**
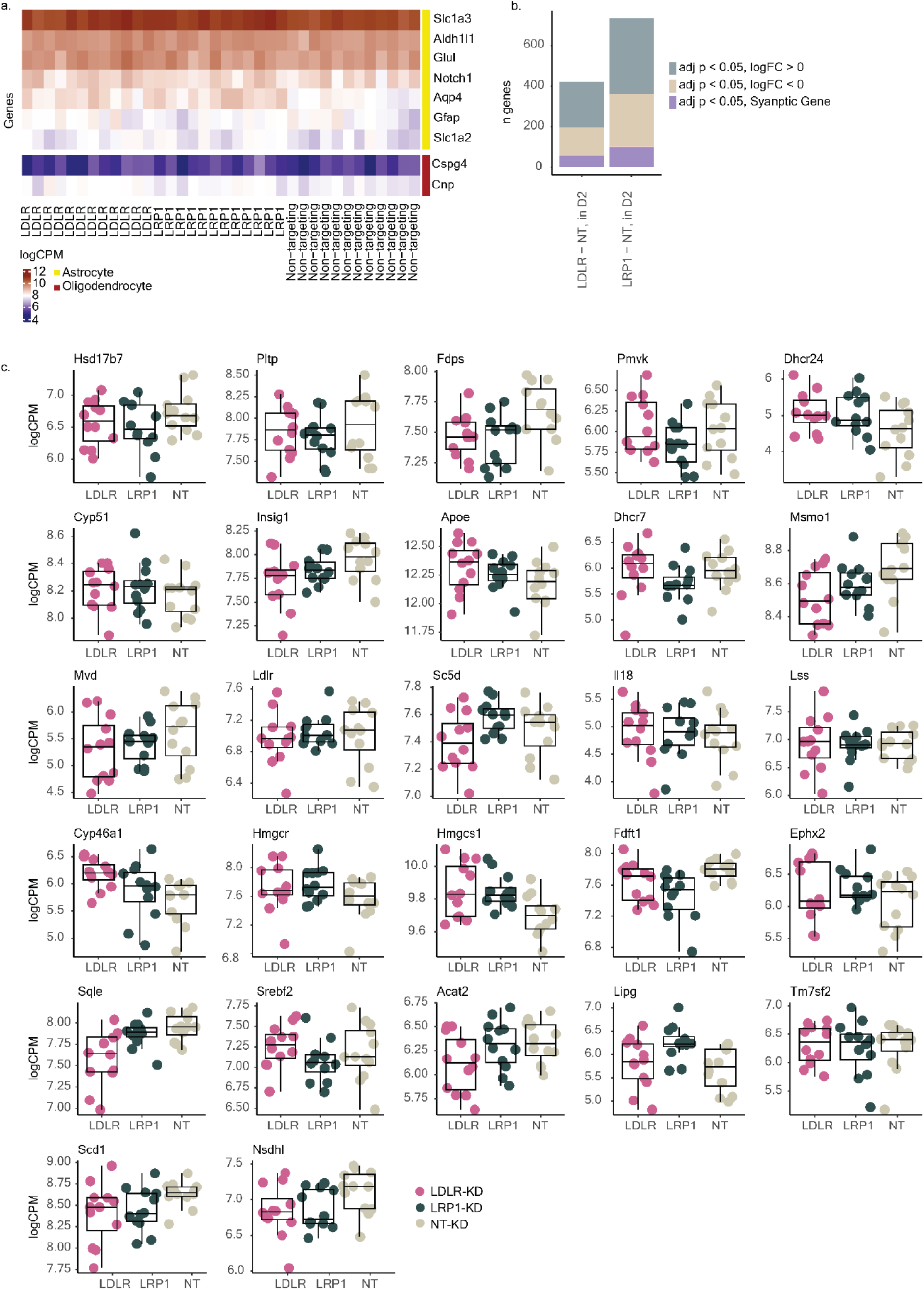
Astrocytes grown with CRISPRi-KD neurons. **a,** Heatmap of normalized expression (logCPM) of canonical astrocyte marker genes in astrocytes grown with *LRP1*-KD and *LDLR*-KD neurons (n = 36). Annotations on the right of the plot show marker gene identity (yellow represents astrocytes, red oligodendrocytes). **b,** Barplot showing the number of DEGs in the astrocytes. Turquoise color shows upregulated genes (adj. p < 0.05 & log2(FC) > 0), beige color shows downregulated genes (adj. p < 0.05 & log2(FC) < 0) and purple color shows the number of significantly DE synaptic genes. **c,** Boxplots of normalized expression (logCPM) for cholesterol biosynthesis genes (n = 27) in astrocytes grown with *LRP1*-KD, *LDLR*-KD and NT-KD neurons. The boxplots’ limits show the upper- and lower quartile ranges, the center lines show the median values. The whiskers show the minimum and maximum values.

**Extended Data Fig. 9:**
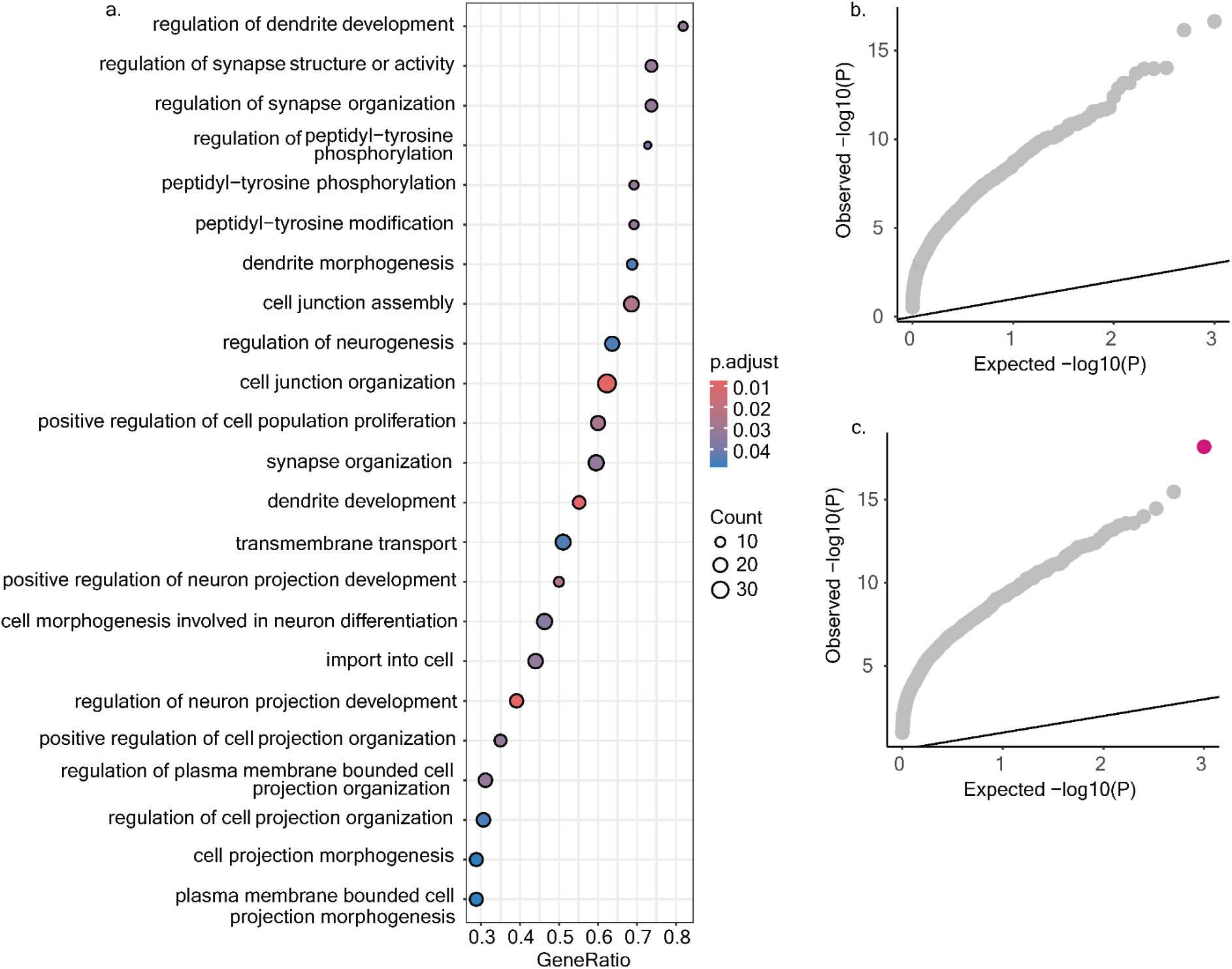
Astrocytes grown with CRISPRi-KD neurons - GO terms and SCHEMA variant burden analysis. **a,** Gene ratio of top enriched GO terms in upregulated genes in astrocytes grown with *LRP1*- and *LDLR*-KD neurons. **b, c,** Qq-plots of p-values from permutation analysis (n = 1,000 permutations) of variant burden in the SCHEMA consortium ^2^ data for downregulated (**b**) and upregulated (**c**) genes in astrocytes grown with *LRP1*- and *LDLR*-KD neurons. The p-values were calculated using Fisher’s exact test. The pink data points highlight enrichment of the gene set in question, the grey points demonstrate the permutation results.

